# Type 2 Diabetes and Obesity Alter Exercise Training-Induced Transcriptional Adaptations to Subcutaneous White Adipose Tissue

**DOI:** 10.1101/2025.10.11.681273

**Authors:** Roeland J.W. Middelbeek, Jiekun Yang, Pasquale Nigro, Benjamin James, Danaé Papadopoulos, Laura K. Simpson, Jeu Park, Maria Vamvini, Li-Lun Ho, Hui Zhang, Nicholas P. Carbone, Michael F. Hirshman, Manolis Kellis, Laurie J. Goodyear

## Abstract

White adipose tissue (WAT) dysfunction contributes to obesity-associated metabolic disease and type 2 diabetes (T2D). Rodent studies have demonstrated that exercise training improves WAT function, but molecular studies investigating exercise training effects on WAT in humans have been limited, particularly in the context of metabolic disease. Here, we defined the subcutaneous WAT (scWAT) transcriptome in middle-aged adults (10 male, 19 female) that were classified by lower BMI (<27 kg/m^2^), higher BMI (≥27 kg/m^2^), and T2D status before and after a 10-week endurance exercise regimen. At baseline, 624 genes were significantly upregulated and 112 genes downregulated in the scWAT from higher BMI participants compared to lower BMI. There was a spectrum of pathway dysregulation in scWAT in higher BMI individuals, ranging from increased markers of extracellular matrix (ECM) deposition and inflammation to decreased circadian rhythm gene expression. In people with T2D, there were additional transcriptomic differences such as translation-related pathways, selenoamino acid metabolism, and proteoglycans. Exercise training had robust effects on the transcriptome regardless of metabolic status, and notably, for the high BMI and T2D groups, training reversed several of the detrimental gene expression patterns in a cell-type-specific manner. These beneficial exercise-induced transcriptomic adaptations significantly correlated with lower levels of free fatty acids and blood pressure, particularly in participants with higher BMI and T2D. By integrating our exercise training-modulated genes with GWAS meta-analysis of physical activity, genes influenced by exercise training in the higher BMI group showed a significant enrichment in genetic associations of exercise traits in the population. A circadian rhythm-related transcription factor NR1D1 was enriched in enhancers linked with both the exercise differentially expressed genes (DEGs) and GWAS signals, suggesting a link between the circadian rhythm and training-induced adaptations. These findings demonstrate that obesity and T2D result in marked, progressive alterations in cell-type specific gene transcription in scWAT, while endurance exercise training reverses many of the metabolic disease-associated adaptations. Identification of novel molecular pathways regulated by exercise training can lead to therapeutic targets for obesity and metabolic disease.

## Introduction

Worldwide rates of obesity have risen dramatically over the last decades, resulting in a corresponding increase in type 2 diabetes (T2D)^1,2^. Obesity is characterized by excessive accumulation of white adipose tissue (WAT), a multi-functional tissue made up of numerous diverse cell types and understanding the molecular makeup of WAT is paramount to deciphering the pathophysiology of obesity and its associated metabolic disorders. WAT is classified based on anatomical location and function, with the two most prominent being subcutaneous (scWAT) and visceral (vWAT), depots that demonstrate significant heterogeneity in cell type composition and metabolic function^3^. In lean healthy individuals, scWAT may be metabolically protective due to increased mitochondrial function, a lower inflammatory state, and the secretion of beneficial adipokines^4^; however, scWAT in people with obesity and T2D have been shown to have detrimental metabolic adaptations including hypertrophy, hypoxia, remodeling and inflammatory macrophage infiltration^5,6^.

Regular exercise is of critical importance in the prevention, treatment, and management of obesity and T2D, in part through improved whole-body glucose homeostasis^7^. Studies investigating rodent models suggest that exercise training provokes significant adaptive responses in scWAT that contribute to the overall metabolic benefits of exercise^8–11^. Using unbiased molecular analyses, we and others have shown that exercise training in mouse models can induce structural and functional adaptations to a number of biological processes and pathways in scWAT including increased innervation and vascularization^12–14^. Moreover, we find that transplantation of scWAT from trained mice into sedentary recipient mice improves glucose homeostasis in recipient mice^7^. While these important findings in rodents suggest a critical role for scWAT in improved metabolic responses with exercise training, the lack of investigation in humans has left a critical knowledge gap in our understanding of how exercise training impacts human scWAT function and gene expression. Of the published human investigations on exercise involving gene profiling, most have focused on specific genes or pathways^15–19^ and/or have only been conducted in healthy^19,20^ or obese males^15^. Given that animal models have shown adipose tissue responses to exercise training are considerably different between females and males^9,21^, it is important to include both women and men in studies of exercise and adipose tissue biology. It is also essential to investigate the effects of exercise in individuals with a range of body mass indexes (BMIs) as well as those with metabolic disease such as T2D. Defining these differences is critical, as it is expected that in 2030, nearly 50% of the US population will be classified as obese^22^.

Here, we determine the effects of obesity with and without T2D on the scWAT transcriptome. We find that in the untrained, baseline condition, scWAT has dramatically different molecular signatures across varying BMI and T2D status. Given that the beneficial health effects of exercise training may differ among individuals with varying metabolic states, we next determined how baseline adiposity influences training-induced molecular adaptations in scWAT. Our results indicate that exercise training has marked beneficial adaptations to scWAT, especially in participants with higher BMI and T2D status, indicating that while exercise benefits all individuals, its impact may be particularly important in these populations. These data provide the foundation for understanding the unique adaptive responses to white adipose tissue that occur with metabolic disease and exercise training in humans.

## Results

### Baseline characteristics

Baseline characteristics of the 29 participants are listed in **Table 1**. Participants consisted of 10 men and 19 women who we classified as lower (<27 kg/m^2^) or higher BMI (≥27 kg/m^2^), resulting in the mean BMI being approximately 6.5 Kg/m^2^ greater in the high BMI group 23.7±2 vs 30.3±3 Kg/m^2^ (full inclusion and exclusion criteria in **Materials and Methods**). As anticipated, higher BMI participants had an overall worse metabolic profile, and of note, cardiorespiratory fitness, as measured by VO_2_peak, was 25% lower in higher BMI participants (**Table 1**). Further subclassification of the subjects showed that 8 of the 16 high BMI subjects had T2D.

**Table 1.** Baseline characteristics of the 29 participants who complete the study. One-way ANOVA was performed to compare these characteristics in the lower and higher BMI groups.

|  | Lower BMI<br>n=13 (3 male/10 female) | Higher BMI<br>n=16 (7 male/9 female) | p-value |
| --- | --- | --- | --- |
| Age (yrs) | 39.5 ± 6.8 | 44.4 ± 6.8 | 0.08 |
| BMI (kg/m<sup>2</sup>) | 23.7 ± 2 | 30.3 ± 3 | **<0.001** |
| Body Fat (%) | 28.5 ± 6.4 | 35.4 ± 8.3 | **0.03** |
| Waist Circ. (cm) | 82.7 ± 5.1 | 99.5 ± 6.4 | **<0.001** |
| FFM (kg) | 47.4 ± 7.5 | 55.6 ± 10.3 | **0.03** |
| Fat Mass (kg) | 19.1 ± 5.3 | 30.3 ± 7.7 | **<0.001** |
| SBP (mmHg) | 116.6 ± 15.7 | 124.3 ± 14.2 | 0.2 |
| DBP (mmHg) | 64.8 ± 10 | 73.6 ± 10.2 | **0.03** |
| RHR (bpm) | 58.7 ± 5.8 | 71.9 ± 10.9 | **<0.001** |
| HbA1C (%) | 5.3 ± .2 | 6.6 ± 1.4 | **0.01** |
| BL Glucose (mmol/l) | 85 ± 5.2 | 110.8 ± 34.5 | **0.02** |
| 2-Hour Glucose (mmol/l) | 106.9 ± 19.5 | 202.5 ± 103.4 | **0.01** |
| FFA (mmol/l) | 0.7 ± 0.2 | 0.5 ± 0.3 | 0.12 |
| Insulin (mU/L) | 3.7 ± 1.2 | 6.2 ± 4.1 | **0.04** |
| AUC | 243.56 ± 27.6 | 381.5 ± 170.8 | **0.01** |
| Creatinine (mg/dl) | 0.8 ± 0.1 | 0.8 ± 0.2 | 0.19 |
| Albumin (g/dl) | 4.2 ± 0.2 | 4.3 ± 0.2 | 0.55 |
| Total Bili (mg/dl) | 0.5 ± 0.2 | 0.5 ± 0.2 | 1.0 |
| Alk Phos (IU/L) | 49 ± 8.2 | 67.4 ± 27.2 | **0.03** |
| AST (IU/L) | 23.5 ± 7.9 | 29 ± 9.4 | 0.12 |
| ALT (IU/L) | 14.7 ± 4.2 | 31.5 ± 15.3 | **<0.001** |
| Cholesterol (mg/dl) | 175.1 ± 38.8 | 184.6 ± 33 | 0.50 |
| Triglycerides (mg/dl) | 75 ± 35.3 | 110.6 ± 47.9 | **0.04** |
| HDL (mg/dl) | 63.3 ± 9.5 | 47.9 ± 11.7 | **<0.001** |
| LDL (mg/dl) | 96.4 ± 33.3 | 114.6 ± 27.8 | 0.14 |
| Total Protein (g/dl) | 6.3 ± 1.8 | 6.4 ± 2.4 | 0.98 |
| Metanephrines (pg/mL) | 34.6 ± 13.2 | 28.8 ± 6.3 | 0.17 |
| Normetanephrines (pg/mL) | 77.3 ± 44 | 65.2 ± 27.6 | 0.44 |
| Total MIE (pg/mL) | 108.4 ± 49.4 | 81.6 ± 28.8 | 0.12 |
| Power (atts) | 185.9 ± 62.1 | 171 ± 59.3 | 0.53 |
| VO<sub>2</sub> (ml/kg/m) | 30.3 ± 8.4 | 22.9 ± 7.1 | **0.02** |

### Pathway analysis demonstrates that lower and higher BMI participants have distinct metabolic pathway regulation in WAT

For our initial analysis comparing baseline parameters between lower and higher BMI, the higher BMI group included both participants with and without T2D. Principal component analysis (PCA) of baseline pre-training samples from the 29 participants showed a separation of lower and higher BMI individuals along the first principal component (PC), which explained 23% of the variance (**Fig. 1A**). When comparing the scWAT from lower and higher BMI groups, the higher BMI group had 624 significantly upregulated and 112 downregulated differentially expressed genes (DEGs) compared to the lower BMI group (**Fig. 1B**). In higher BMI individuals, of the top 10 upregulated genes determined by P-value, three are related to the extracellular matrix (ECM; *EMILIN2*, *CCN5* and *LTBP2*), six are involved in leukocyte migration regulation or are leukocyte markers (*MPEG1*, *CYBB*, *EMILIN2*, *AIF1*, *CCL2* and *PTPRC*), and two genes (*MPEG1* and *AIF1)* are linked to macrophage activation^23^ and increased inflammation in obesity^24^ (**Fig. 1B, Table S2**).

**Figure 1.**
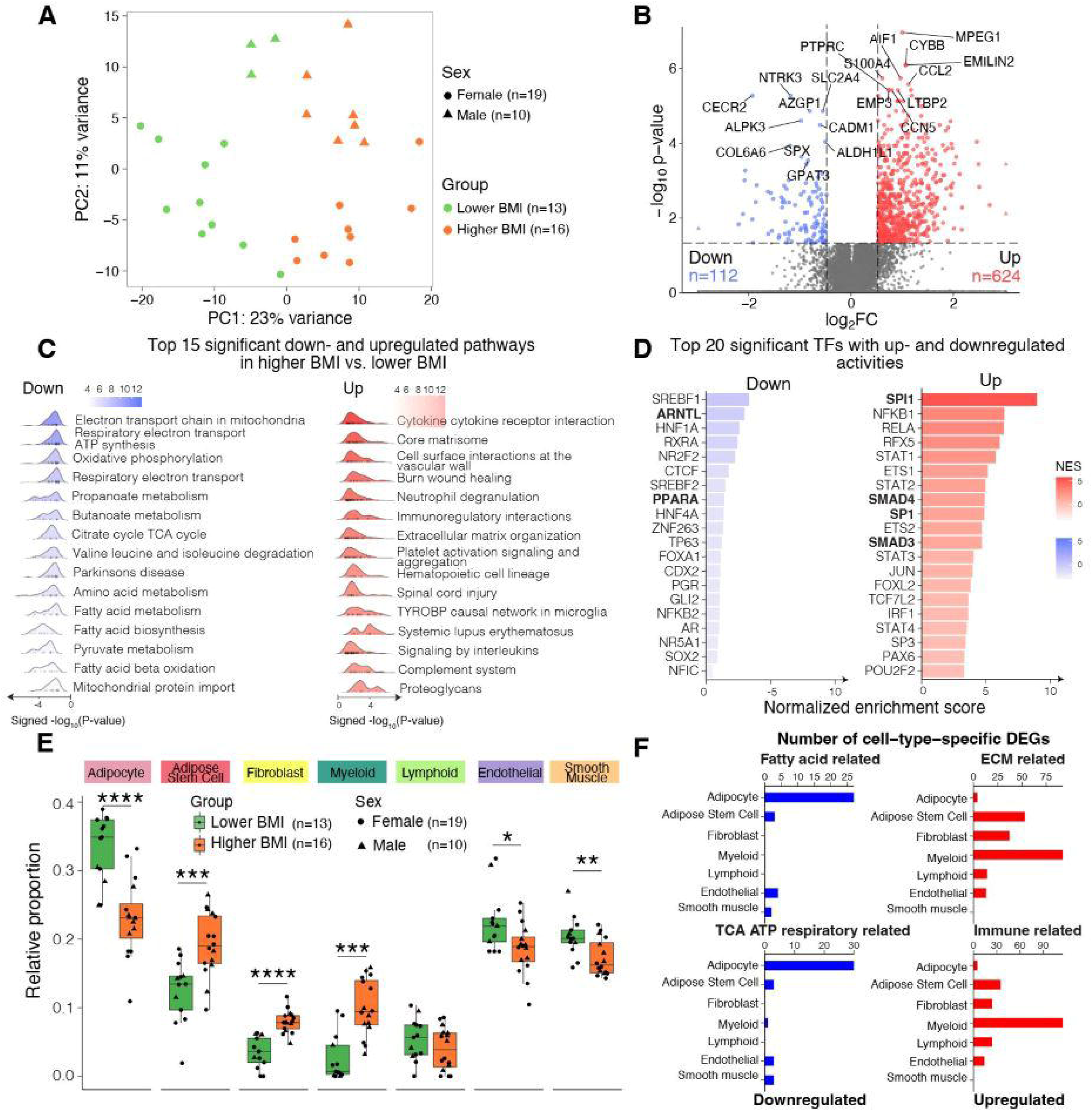
Distinct genes, pathways and cell type proportions in scWAT between lower and higher BMI subjects at baseline. **A**, PCA plot for baseline RNA-seq data from the 29 subjects included in this study. The plot is colored by lower or higher BMI. **B**, Volcano plot showing the differentially expressed genes (DEGs) between the higher and lower BMI groups, with the top 10 significantly up- and downregulated genes labeled. **C**, Ridge plot showing the top 15 significant pathways enriched by the up- and downregulated DEGs. **D**, Bar plots showing the inferred transcription factors (TFs) enriched in the up- and downregulated DEGs. TFs discussed in the text are in bold. **E**, Box plots comparing the deconvolved cell type proportions between the lower and higher BMI groups. The boxplot visualizes the median, the first and third quartiles and 1.5*inter-quartile range from the hinge. *, p<0.05; **, p<0.01; ***, p<0.001; ****, p<0.0001. **F**, Bar plots showing the number of cell-type-specific DEGs in the four pathway groups in higher BMI vs. lower BMI. Up- and down-regulated pathway groups are colored in red and blue, respectively.

Next, we performed pathway analysis on the ranked full gene list using fgsea^25^. Consistent with the functions of the top upregulated genes, the top 15 pathways upregulated in higher BMI participants at baseline concentrate on two biological pathway groups: 1) the immune pathway group including cytokine interaction, neutrophil degranulation, immunoregulatory interaction, complement system, and interleukin signaling; and 2) the ECM pathway group, including ECM organization, proteoglycans, and the core matrisome (**Fig. 1C, Table S3**). Key metabolic pathways significantly downregulated in higher BMI participants include pathways related to the TCA cycle, mitochondrial function, oxidative phosphorylation, amino acid metabolism, and fatty acid biosynthesis and metabolism (**Fig. 1C, Table S3**). Taken together, these results demonstrate that higher BMI individuals have upregulation of inflammation-related and ECM pathway transcriptional signatures, and downregulation of key metabolic pathways, a distinct gene expression profile which may predispose higher BMI individuals to scWAT adaptations that lead to adipose tissue dysfunction.

To determine potential master regulators for the DEGs and enriched pathways, we performed transcription factor (TF) enrichment analysis using DoRothEA^26^. Only the TF-target interactions supported by different lines of evidence (literature-curated resources, ChIP-seq peaks, TF binding motifs on promoters, and inference from gene expression) were included in this analysis. Higher BMI participants showed increased activities of TFs regulating inflammatory transcriptional programs in macrophages (SPI1^27^), and ECM deposition (SMAD3-SMAD4-SP1 complex^28^; **Fig. 1D, Table S4**). Notable TFs decreased in higher BMI participants include ARNTL/BMAL1, a core molecular clock component with adipocyte-specific deletion resulting in obesity^29^ and PPAR⍺, a key gene involved in amino acid and lipid metabolism in WAT^30^(**Fig. 1D, Table S4**). To understand the contribution of sexual dimorphism in adipose tissue in these results, we repeated the same set of analyses (DEG, pathway, and TF) for male and female participants separately and observed significantly more DEGs in females than in males (**Fig. S1A,B; Table S2).** Females also showed pathways and master regulators enrichment similar to the full combined analysis (**Fig. S1C-F; Table S3,4**).

### BMI-linked variations in scWAT cell-type composition

WAT is a heterogeneous tissue and contains numerous cell types including adipocytes, immune cells (i.e. myeloid and lymphoid cells), adipose stem cells (ASCs), fibroblasts, and vascular-related cells (endothelial and smooth muscle cells)^13,31^. To understand if there are differences in cell type proportions and cell type-specific transcriptional profiles between lower BMI and higher BMI participants, we deconvolved the bulk RNA-seq data into 7 major cell types by referencing two publicly available single-cell datasets^32,33^. Compared to lower BMI participants, higher BMI participants had significantly lower percentages of adipocytes, endothelial, and smooth muscle cells, while there were significantly higher percentages of ASCs, fibroblasts, and myeloid cells (**Fig. 1E**). The effects of higher BMI on cell type proportion went in the same direction in males and females (**Fig. S2A**). Interestingly, there were no differences between male and female participants in cell type proportion in any of the cell types for both lower BMI and higher BMI, except for mature adipocytes in lower BMI, which showed higher proportions in females than in males (**Fig. S2B**).

To identify the specific cell types accounting for the major pathway differences in higher vs. lower BMI we compared cell type-specific gene expression matrices (**Fig. 1F**). Decreases in fatty acid metabolism and TCA cycle pathway related genes in the higher BMI subjects occurred predominantly in adipocytes, whereas increases in the ECM and immune pathway related genes in the higher BMI participants were greatest in myeloid cells and ASCs. These results demonstrate that the deconvolved gene expression differences that occur in higher vs. lower BMI participants, and in participants with T2D, manifest in distinct cell types.

### Comparison of scWAT gene profile in high BMI participants with and without T2D

A series of analyses were performed to determine if there are transcriptional differences in the higher BMI group with T2D compared to the higher BMI group without T2D. The higher BMI participants with T2D are referred to as “T2D” in the current and following sections for simplicity. Re-coloring the PCA plot revealed a gradual separation of these two groups along PC1 (**Fig. 2A**). Higher BMI participants without T2D lie between lower BMI and T2D participants, demonstrating the progressive phenotype trajectory from lower BMI to higher BMI, to higher BMI with T2D (**Fig. 2A**).

**Figure 2.**
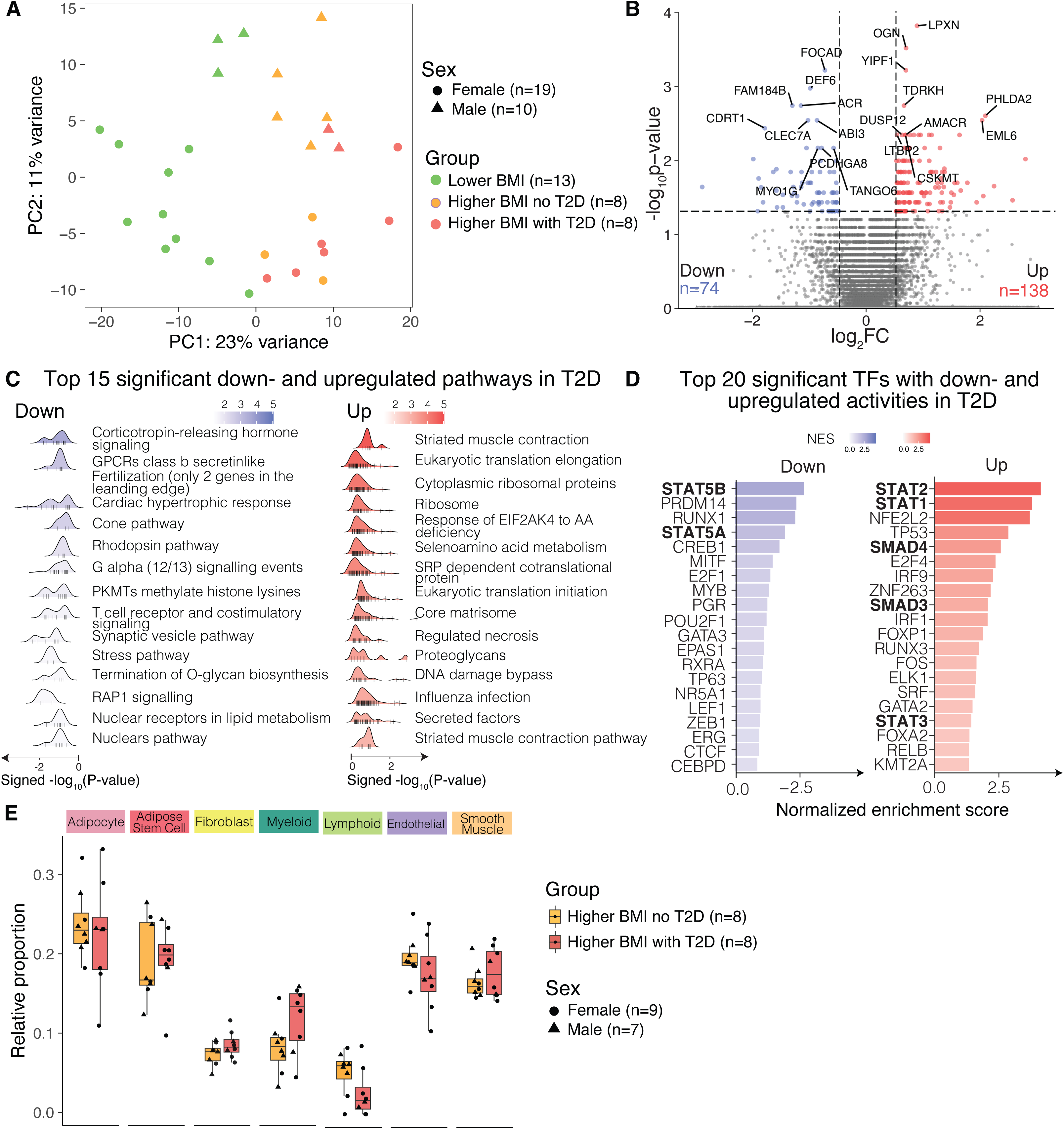
Distinct genes and pathways in scWAT between higher BMI subjects with and without T2D at baseline. **A**, PCA plot for baseline RNA-seq data from the 29 subjects included in this study. The plot is colored by lower, higher BMI no T2D, and higher BMI with T2D. **B**, Volcano plot showing the differentially expressed genes (DEGs) between the higher BMI group with T2D vs. without, with the top 10 significantly up- and downregulated genes labeled. **C**, Ridge plot showing the top 15 significant pathways enriched by the up- and downregulated DEGs in the higher BMI group with T2D vs. without. **D**, Bar plots showing the inferred transcription factors (TFs) enriched in the up- and down-regulated DEGs in the higher BMI group with T2D vs. without. TFs discussed in the text are in bold. **E**, Box plots comparing the deconvolved cell type proportions between the two higher BMI groups: with and without T2D. The boxplot visualizes the median, the first and third quartiles and 1.5*inter-quartile range from the hinge.

For DEGs, there were 138 upregulated genes in T2D individuals compared to higher BMI without T2D (**Fig. 2B, Table S2**). These upregulated genes were enriched in ribosomal and translation related pathways, selenoamino acid metabolism, core matrisome, regulated necrosis, proteoglycans, and secreted factors (**Fig. 2C, Table S3**). T2D participants had 74 downregulated genes, including genes involved in the innate immune system (e.g. *CLEC7A)* (**Fig. 2B, Table S2**), and pathways enriched in corticotropin-releasing hormone signaling, T cell receptor and costimulatory signaling, and synaptic vesicle pathways (**Fig. 2C, Table S3**).

TF analysis showed increased target gene expression levels of TFs belonging to the STAT family (STAT2, STAT1, and STAT3) and SMAD family (SMAD4, SMAD3) in T2D vs. higher BMI participants (**Fig. 2D; Fig. S3E,F; Table S4**). SMAD proteins have been implicated in increased adiposity, insulin resistance, and T2D^34^, while the STAT family of TFs are activated by cytokine signaling, and involved in immune responses^35^. The upregulated TF STAT1 has been linked to adipose tissue inflammation^36^. We found that STAT5A and STAT5B were downregulated in T2D. STAT5A and STAT5B mediate lipid mobilization from adipocytes^37^, and studies in male mice lacking both STAT5A/B proteins in mature adipocytes exhibit impaired WAT lipolysis^38^.

Importantly, we did not observe any difference in cell type proportions between T2D and higher BMI without T2D participants (**Fig. 2E**), demonstrating that the changes in T2D cell phenotype are not due to alterations in cell type proportions. Another interesting finding was that when we compared the three groups separately (lower BMI vs higher BMI without T2D vsT2D), there was a gradual increase in expression of ECM-related pathways (**Fig. S3C,D; Table S3**), suggesting increased ECM remodeling and WAT tissue fibrosis occurs as a continuum with increasing metabolic disease.

### Exercise training improves cardiorespiratory fitness in participants without T2D, and glycemic control in T2D participants

Following baseline assessments, participants completed a 10-week exercise training program. As described in Materials and Methods, exercise training in lower BMI participants was done by either moderate intensity training (MIT) or high intensity interval training (HIIT). Both training modalities induced a similar relative increase in VO_2_peak and Peak Power (10 and 11%, respectively, p=0.81, **Fig. S4A**). In line with similar responses in VO_2_peak with training, there was not a significant difference in scWAT transcriptomic change between the two training types in the lower BMI group, and therefore, for subsequent analysis these two training groups were combined.

Exercise training increased VO_2_peak (**Fig. 3A**) and peak power (**Fig. 3B**) in lower and higher BMI participants without T2D, but training did not improve these parameters in T2D participants, despite all three groups successfully completing the training program. Participants who had lower BMI and higher BMI without T2D had normal glucose tolerance (**Fig. 3C,D**) and HbA1c% (**Fig. 3F**) at baseline and exercise training did not change these parameters. Although T2D participants did not increase VO_2_peak or power, the exercise training intervention improved glycemic control, as evidenced by an improved glucose tolerance as assessed by the glucose excursion curve (**Fig. 3E**), glucose area under the curve (AUC) (**Fig. 3F**), and improved HBA1C% (**Fig. 3G**). There was no change in insulin AUC during the OGTT (**Fig. 3H**). Exercise training did not alter body weight, fat mass, and lean mass in any of the groups (**Fig. 3I-K**), allowing us to specifically assess the effects of the exercise intervention.

**Figure 3.**
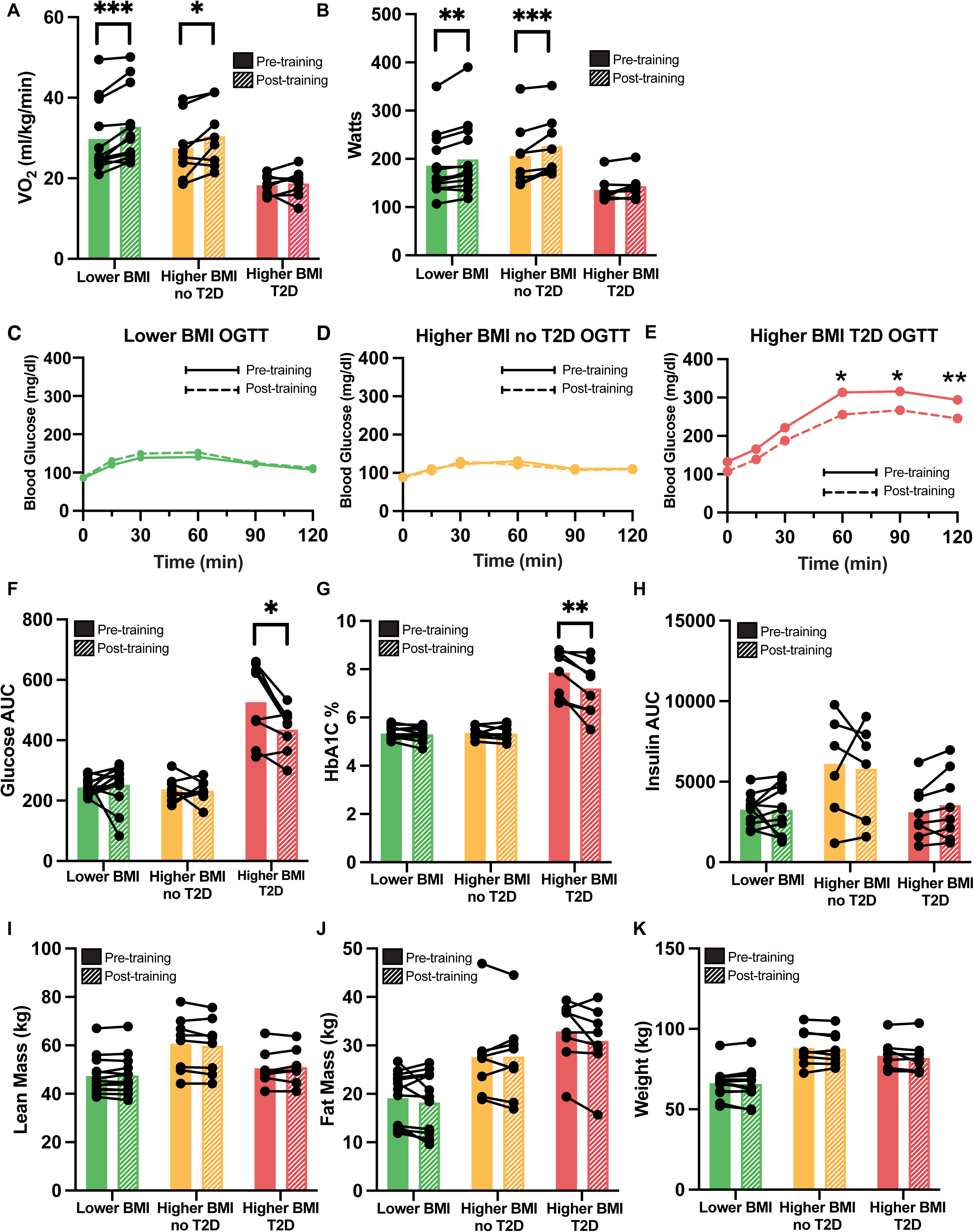
Effects of exercise training on glucose tolerance and body composition in all groups (Lower BMI n=13; Higher BMI without T2D n=8; Higher BMI with T2D n=8). **A**, Bar graph showing VOL peak before and after training in each group. **B**, Bar graph showing peak watts achieved during VOL peak before and after training in each group. **C,D,E** Line graph showing the plasma glucose concentration during oral glucose tolerance test (OGTT) before and after training in the lower BMI group (**C**), higher BMI group without T2D (**D**), and higher BMI group with T2D (**E**). F,G,H Bar graph showing area under the curve (AUC) of plasma glucose (**F**), HbA1c%(**G**), and area under the curve (AUC) of insulin during OGTT (**H**), before and after training in each group. **I,J,K** Bar graph showing lean mass (**I**), fat mass (**J**), and body weight (**K**), before and after training in each group.

### Exercise training-induced reversal of gene expression alterations in higher BMI and T2D participants

We first analyzed the effects of exercise training by comparing the lower BMI group with the combined higher BMI group. Exercise training induced transcriptomic changes in scWAT, as shown by differences in pre- and post-training samples based on PC euclidean distances in lower and higher BMI participants (**Fig. 4A**). Exercise training in lower BMI participants increased expression of 228 genes and downregulated only 19 genes (**Fig. 4B and listed in Table S5**) whereas training in the higher BMI group increased 76 genes and downregulated 46 genes (**Fig. 4C, Table S5**). Surprisingly, the training-induced changes in specific scWAT gene expression were strikingly different in comparing lower BMI and higher BMI participants (**Fig. 4D**). In fact, training in lower and higher BMI participants had only three commonly upregulated genes (upper right quadrant; *IQUB*, *TOR2A*, and *KLHL26)* and only one commonly downregulated gene (lower left quadrant; *CRISPLD1)*. One gene, *SELL*, was upregulated by training in the lower BMI group but downregulated in the higher BMI group (upper left quadrant). The limited number of overlapping DEGs in the two groups show that exercise training has distinct effects on scWAT gene expression in lower vs. higher BMI participants.

**Figure 4.**
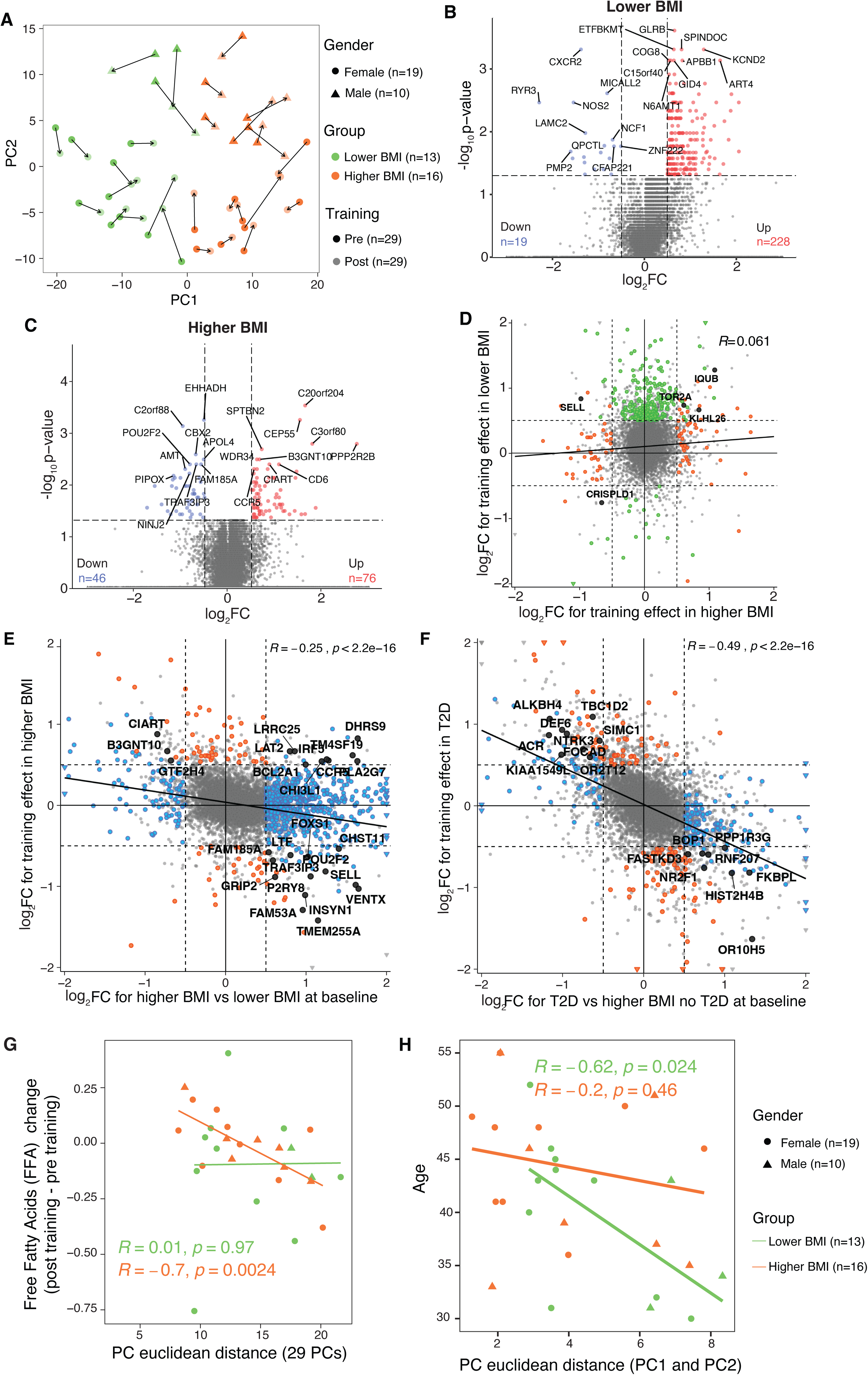
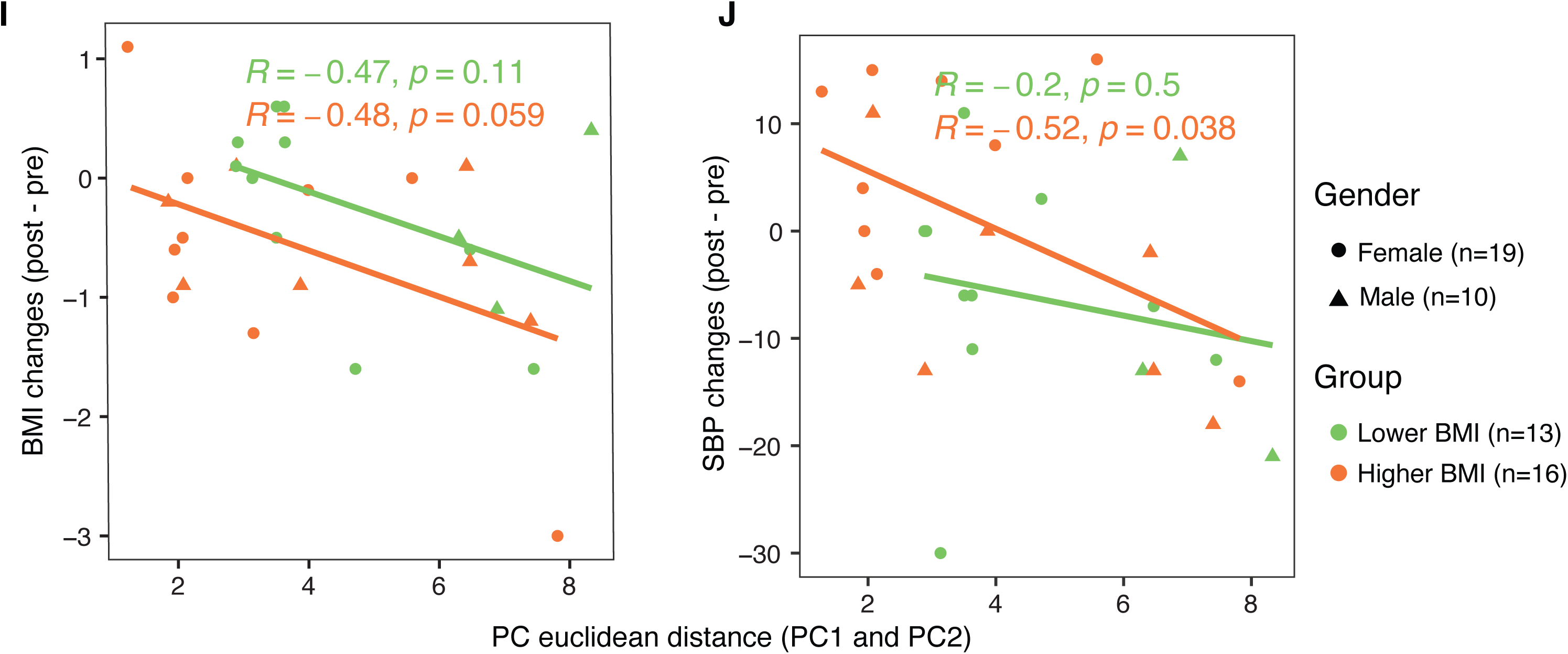
Exercise training-induced global transcriptional shifts and gene-level reversals. **A**, PCA plot for pairwise RNA-seq data from the 29 subjects included in this study before and after exercise training. Arrows point from the pre-training sample to the post training sample for each subject. **B,C** Volcano plot showing the differentially expressed genes (DEGs) post-training compared to pre-training in the lower BMI (**B**) and higher BMI group (**C**), with the top 10 significantly up- and downregulated genes labeled. **D**, Scatter plot showing logL fold changes in training-induced gene expression in higher BMI (x-axis) vs. lower BMI (y-axis); green, orange, black, and gray dots indicate significance in lower BMI only, higher BMI only, both, and neither, respectively*, p<0.05 E,F, Scatterplot comparing the log2 fold changes for the training effect in higher BMI vs. BMI effect at baseline (**E**) or the log2 fold changes for the training effect in T2D vs. T2D effect at baseline (**F**). Black dots indicate genes that are significant in both comparisons: baseline expression between higher BMI vs. lower BMI (x-axis) and expression changes with training (y-axis). Blue dots indicate genes that are significant only in the baseline comparison between higher BMI and lower BMI, and orange dots indicate genes that are significant only in the training-induced changes within the higher BMI group. Grey dots represent genes that were not statistically significant in either comparison. **G**, Scatterplot correlating PC Euclidean distance, representing global transcriptional shifts, with free fatty acids change post-training compared to pre-training for the lower and higher BMI groups separately. Pearson correlation coefficient ® and p value are labeled on the plot. **H,I,J** Scatterplot showing correlation between age (**H**), BMI change (**I**), SBP change (**J**), and degree of scWAT transcriptomic changes across all groups.

To assess the transcriptome-wide impact of exercise training on obesity- and diabetes-related gene expression, we performed correlation analyses between baseline expression profiles and training-induced changes (**Fig. 4E and 4F**). A highly significant negative correlation was observed between BMI-associated gene expression and the transcriptional response to exercise (**Fig. 4E**), indicating that genes upregulated in obesity tended to be downregulated following training. This trend was even more pronounced for genes elevated in T2D, which showed a stronger inverse correlation with the exercise response (**Fig. 4F**), suggesting that exercise counteracts both obesity and T2D-associated transcriptional programs. As the higher BMI group with T2D showed significant improvement in glucose metabolism post-training, we analyzed the higher BMI group by T2D status and found that in the T2D group, exercise training uniquely regulated 17 genes that showed opposite direction of gene expression post-training compared to baseline (**Fig. 4F**). Several of these genes are linked to glucose metabolism (i.e. PPP1R3G, FKBPL, NR2F1, and BOP1), suggesting their potential contribution to the improved glycemic control observed after training in the T2D group.

### Transcriptional shifts in scWAT induced by exercise training and their clinical correlations

By performing regression analysis with the clinical variables, we identified the scWAT transcriptomic changes that correlated with key cardiometabolic factors. Interestingly, global transcriptomic adaptations post-exercise training inversely correlated with changes in free fatty acids (FFAs) in the higher BMI group, but not in the lower BMI group (**Fig. 4G**). Besides FFAs, age showed an inverse correlation with scWAT transcriptomic changes in lower BMI individuals only (**Fig. 4H**). Despite no significant reduction in BMI post-training, changes in BMI showed a trend (p=0.059) to be inversely correlated with scWAT transcriptomic changes in the higher BMI group (**Fig. 4I**), meaning more scWAT transcriptomic changes tended to correspond to larger BMI reductions for participants with higher baseline BMI. Additionally, there was a significant negative correlation between systolic blood pressure (SBP) changes and the scWAT transcriptomic changes only in the higher BMI group (**Fig. 4J**), which suggests a connection between exercise training-induced scWAT remodeling and improved blood pressure. Although causation cannot be established, exercise training-induced transcriptomic changes in the higher BMI group correlated with FFAs, SBP, and trended with BMI. Taken together, these correlation data highlight the potential involvement of scWAT transcriptomic changes in the significant phenotypic adaptations post-training in higher BMI individuals.

### Training induced changes in transcription factors across BMI groups and T2D status

To understand transcriptional responses to exercise training, we next investigated transcription factor (TF) expression. Exercise induced three pathways across BMI groups, with one prime example being upregulation of pathways targeting protein synthesis (**Fig. 5A,B, Table S6**). Nine of the top 20 upregulated TFs were shared (**Fig. 5C,D**; shared TFs in blue bold). Among the training-induced upregulated TFs, the androgen receptor (AR) has shown a significant role in modulating the scWAT adaptation to exercise training, including induction of lipolysis^9^. Four TFs were commonly downregulated (**Fig. 5C,D**; shared TFs in blue bold), including the key diabetes-risk genes HNF1A, associated with monogenic (MODY3) diabetes^39^, and TCF7L2, the most potent locus for T2D risk with fundamental developmental and metabolic roles in adipose tissue^40^. While these regulated TFs point to common regulatory mechanisms, exercise training also induced unique transcriptional patterns specific to each group.

**Figure 5.**
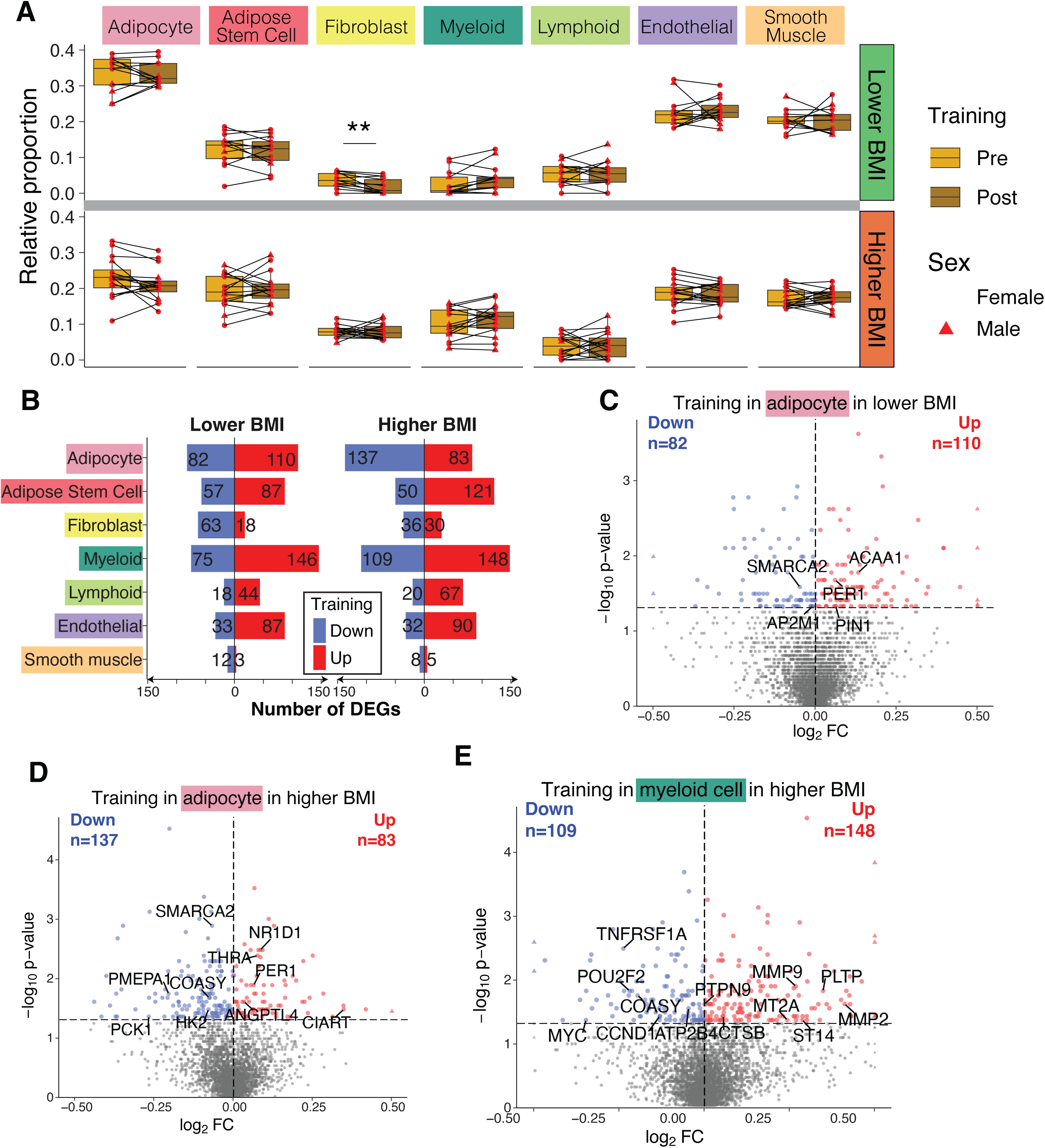
Distinct pathways and transcription factors in scWAT associated with training in lower and higher BMI subjects. **A**, Ridge plot showing the top 15 significant pathways enriched by the up- and downregulated DEGs in the lower BMI group. **B**, Ridge plot showing the top 15 significant pathways enriched by the up- and downregulated DEGs in the higher BMI group. **C**, Bar plot showing the inferred transcription factors (TFs) enriched in the up- and downregulated DEGs in the lower BMI group, expressed as Normalized Enrichment Score (NES). **D**, Bar plot showing the inferred TFs enriched in the up- and downregulated DEGs in the higher BMI group. **E**, Bar plots showing the inferred TFs enriched in the up- and downregulated DEGs in the lower BMI MIT group. **F**, Bar plot showing the inferred TFs enriched in the up- and downregulated DEGs in the lower BMI HIT group. **G**, Bar plot showing the inferred TFs enriched in the up- and downregulated DEGs in the higher BMI no T2D group. **H**, Bar plot showing the inferred TFs enriched in the up- and downregulated DEGs in the higher BMI T2D group.

Lower BMI participants uniquely upregulated PPAR signaling and oxidative phosphorylation with training (**Fig. 5A**). PPARG, as a top TF upregulated by exercise training only in the lower BMI group, is a core component of the PPAR signaling pathway.(**Fig. 5C; bolded in black**). Exercise training in higher BMI participants downregulated key metabolic pathways, including the TCA cycle, glycolysis, and fatty acid biosynthesis (**Fig. 5B**) which may indicate reduced storage of fatty acids in WAT. The TF ARNTL/BMAL1 **(Fig. 5D, bolded in black)**, which is a key factor in circadian rhythm regulation and is downregulated at baseline in individuals with higher BMI, showed increased expression of its target genes following exercise training in participants with higher BMI (**Fig. 1D, 5D**), suggesting an interplay between circadian rhythm, obesity, and the effects of exercise training.

These distinctions were further sharpened by T2D status. Specifically, the upregulation of ARNTL following exercise occurred only in higher BMI participants without T2D, indicating a blunted adaptive response in those with T2D (**Fig. 5E,F; Table S7**). Individuals with T2D uniquely upregulated other TFs, such as POU2F1/OCT-1, a transcription factor that has been associated with increased metformin action in people with obesity^41^. Taken together, exercise training affects numerous TFs in a manner dependent on both BMI and T2D status, highlighting that these distinct responses may contribute to specific metabolic improvements post-training.

One of the most consistent effects of exercise training on TF expression is the upregulation of genes involved in extracellular matrix (ECM) remodeling (**Fig. S6A,B**). This response was more pronounced in participants with lower BMI, who exhibited a greater number of differentially expressed genes associated with elastic fiber-related ECM remodeling compared to those with higher BMI (**Fig. S6C,D**). In addition to ECM-related changes, exercise strongly influenced genes within the PPAR signaling pathway in lower BMI individuals (**Table S7**). Specifically, MMP9, a matrix metalloproteinase and known PPARγ target, was upregulated by training in the lower BMI group (**Fig. S6A**), demonstrating a link between PPAR signaling and ECM remodeling in response to exercise. TF gene expression patterns also differed by sex and BMI group (**Fig. S6G–J**). One notable example is DBP, a transcriptional activator and known target of the core clock gene ARNTL/BMAL1. DBP exhibited sex-specific regulation, showing increased expression only in lower BMI males following training (**Fig. S6I**). This pattern is consistent with our previous findings in mice, where a homologous gene (*Dbp*) was significantly upregulated by exercise training, supporting a conserved cross-species effect¹³.

### Exercise training reprograms cell-type composition and gene expression in scWAT, with distinct patterns shaped by BMI and sex

A paired analysis of the deconvolved cell type proportions post exercise-training revealed a significant decrease in fibroblasts in lower BMI participants which may reflect enhanced tissue plasticity **(Fig. 6A, S6A)**. The reduced proportion of fibroblasts may reflect higher degrees of tissue remodeling and plasticity after exercise training in lower BMI participants. In contrast, fibroblast proportions were unchanged by training in the higher BMI group, potentially reflecting pre-existing fibrosis and impaired remodeling linked to metabolic dysfunction^42^. Within the endothelial compartment, sex-specific effects were noted. Lower BMI females exhibited a significant increase in endothelial cell proportions post-training—a pattern not observed in their male counterparts **(Fig. S6A)**. Higher BMI males showed a reduction in mature adipocyte proportions after exercise training, suggesting altered lipid storage dynamics unique to this participant subgroup. Together, these findings highlight BMI-dependent variation in cell-type-specific responses to exercise training, with additional modulation by sex.

**Figure 6.**
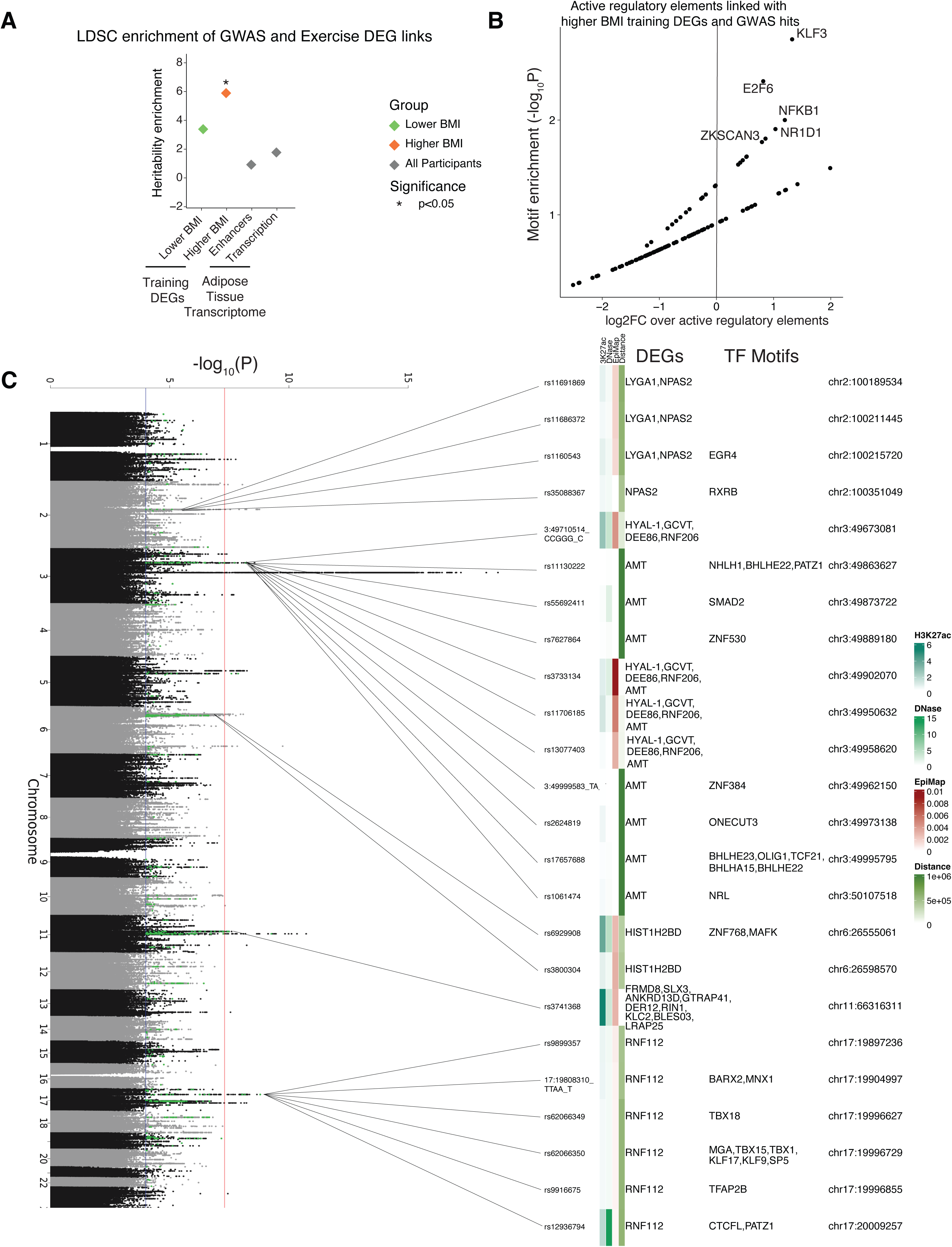
Cell-type-specific training effects in lower and higher BMI groups. **A**, Box plots showing deconvolved cell type proportions in scWAT before and after exercise training in lower and higher BMI groups. The boxplot visualizes the median, the first and third quartiles and 1.5*inter-quartile range from the hinge. Paired Wilcoxon Rank Sum test was performed. *, p<0.05. **B**, Bar plots representing the number of up-(in red) and down-(in blue) regulated cell-type-specific DEGs in the lower and higher BMI groups. **C**,**D** Volcano plots showing training-induced DEGs in adipocytes in lower BMI group (**C**) and in higher BMI group (**D**). **E**, Volcano plot showing training-induced DEGs in myeloid cells in higher BMI group.

Next, DEG responses to exercise training were examined across cell types **(Fig. 6B)**. Mature adipocytes, adipose stem cells (ASCs), and myeloid cells exhibited the highest number of DEGs following training, underscoring their central role in scWAT adaptation. Notably, higher BMI participants demonstrated stronger downregulation of adipocyte and myeloid gene programs with training compared to those with lower BMI, suggesting a heightened transcriptional response. Examples of cell-specific gene regulation include circadian rhythm components, extracellular matrix (ECM)-related genes, and immune response markers **(Fig. 6C,D,E)**. In mature adipocytes, PER1, NR1D1 (Rev-Erbα), and CIART were upregulated post-training **(Fig. 6C,D)**. Within the myeloid lineage, exercise training stimulated ECM-degrading enzymes—MMP9 in higher BMI participants and CTSL in lower BMI participants—suggesting distinct remodeling trajectories (**Fig. 6E, S6B**).

### Exercise training-induced DEGs in scWAT showed enrichment in genetic associations linked to exercise traits

Although traits for physical activity and sedentary behavior are only moderately heritable, genetics of these traits could inform the biological mechanisms underlying these modifiable lifestyle determinants^43^. To elucidate genomic mechanisms and link genes with genetic variants, we determined which exercise training-induced DEGs identified in the current study are enriched in single nucleotide polymorphisms (SNPs) implicated in genome-wide association studies (GWASs) for exercise training-related traits. We also determined which DEGs are located adjacent to scWAT-specific regulatory elements and scWAT-specific expression Quantitative Trait Loci (eQTL). To achieve this goal, we leveraged 1) a multi-ancestry meta-analysis of GWASs for self-reported physical activity and sedentary behaviors, which included up to 703,901 individuals^44^; 2) 2.2M candidate cis-Regulatory Elements (cCREs) from the ENCODE consortium; 3) 2.93M scWAT-specific eQTLs across 581 individuals as reported by GTEx (version 8); and 4) stratified linkage disequilibrium (LD) score regression, which estimates heritability and genetic correlation from GWAS summary statistics^45,46^. By integrating our exercise-modulated genes with the GWAS meta-analysis variants associated with moderate-to-vigorous intensity physical activity during leisure time (GCST90104341), we observed significant enrichment of genetic associations linked to exercise traits in exercise training-induced DEGs in the higher BMI group (**Fig. 7A**). We used regulatory elements linked via proximity, eQTLs, and correlation to nominate for each group of DEGs possible GWAS variants. The exercise-modulated genes with genetic associations linked to exercise in all the participant groups except for the higher BMI male group showed higher genetic enrichment compared to the aggregated scWAT transcription or enhancer activity, suggesting these scWAT genes may play a causal role in determining physical activity levels (**Fig. 7A**).

To identify potential TFs driving putative causal gene expression changes, we performed TF enrichment analysis against all the ENCODE cCREs. KLF3, E2F6, NFKB1, NR1D1 and ZKSCAN3 were the top five enriched TFs in the active regulatory elements that are functionally implicated with the training DEGs in the higher BMI group and the GWAS hits for physical activity (**Fig. 7B**). NR1D1 was one of the genes that we observed significantly increased expression in mature adipocytes with exercise training (**Fig. 6D**). For the other participant groups, ZEB1 was significantly enriched in the females with lower and higher BMI, consistent with our previous RNA only analysis results (**Fig. S7A,B, Fig. 5C,D**). TFAP2A, and TFAP4, part of the TFAP family of genes, have been implicated in cell proliferation and as regulators of lipid droplet biogenesis^47^. These genes were significantly enriched in the lower BMI, and higher BMI males (TFAP2), and higher BMI female groups (TFAP4), suggesting a potential role for the TFAP family in regulating white adipose tissue adaptations post exercise training in higher BMI (**Fig. S7B,C, Fig. 5C,D**). To identify causal genes mediating exercise training effects in scWAT, we visualized all the GWAS variants linked with the exercise training-induced DEGs using a Manhattan plot (**Fig. 7C**), demonstrating numerous DEGs and their TF motifs, including TFAP2B. Taken together, this analysis uncovers potential genetic mediators underlying the effects of exercise training on scWAT and their link to physical activity levels, and in doing so, identifies potential targets for future mechanistic studies.

## Discussion

Obesity-driven pathological changes in WAT promote the development of metabolic disease, and here, we identify a progressive change in scWAT gene expression with increasing adiposity and T2D that likely contribute to this dysregulation. In contrast to obesity, regular exercise can prevent obesity-associated diseases and reverse the progression to T2D, effects that occur through numerous physiological^48,49^ and presumably transcriptional adaptations to multiple tissues. Although there had been a gap in our knowledge of the molecular underpinnings that occur in scWAT in response to exercise training in people with metabolic disease, in the current study we define the scWAT transcriptome and determine that training results in robust changes in gene expression in people with a range of BMI and T2D status. Exercise training decreases numerous genes that were upregulated with obesity and T2D and alters cell type-specific proportions. These exercise-induced transcriptional shifts were further connected to scWAT-specific distal enhancers and genetic variants (SNPs) associated with physical activity, based on genome-wide association studies (GWAS). Together, these four layers of analysis—baseline BMI differences, exercise-induced transcriptomic adaptations, cell type-specific changes, and genetic regulatory links—provide a foundation for uncovering the molecular mechanisms through which exercise training benefits metabolic health in human scWAT.

Our integrative analysis suggests that exercise training-induced transcriptional changes in scWAT are not only responsive to environmental stimuli but are also enriched in genetic variants associated with physical activity traits. This finding highlights a potential bidirectional relationship: while exercise remodels the transcriptomic landscape of adipose tissue, inherited genetic variation may predispose individuals to differential responsiveness to exercise. In particular, enrichment of GWAS variants in DEGs from the higher BMI group indicates that genetic architecture may contribute to variability in exercise adaptation in individuals with higher BMI. The identification of TFs such as NR1D1, ZEB1, and members of the TFAP family further underscore the possibility that specific regulatory networks integrate genetic predisposition with exercise-induced signaling to drive adipose tissue remodeling. Although these analyses do not establish causality, the identified genes and TFs represent important targets for future mechanistic studies. Together, these findings link genetic determinants of physical activity with scWAT transcriptional remodeling, suggesting that host genetics may shape the systemic benefits of exercise.

Prior to exercise training there were greater markers of ECM deposition in higher BMI and T2D participants, suggesting that a thicker ECM structure may underlie functional and structural differences in scWAT associated with obesity and T2D, potentially as a consequence of long-term low-grade inflammation and subsequent tissue remodeling. Opposite to the effects of obesity, in all participants exercise training upregulated genes that encode enzymes that degrade ECM suggesting remodeling of ECM, consistent with our work in mice showing decreased scWAT ECM structure following voluntary wheel running exercise training^13,14^. By deconvolving human scWAT composition, we observed that ASCs and myeloid cells are important players mediating ECM changes in scWAT, as key ECM DEGs were expressed in these cell types, and genes related to ECM degradation were upregulated post exercise-training. Taken together, these observations suggest that exercise training induces ECM degradation pathways in WAT, thereby reversing some of the detrimental adaptations of long-term obesity to ECM in scWAT in humans.

Analysis of scWAT composition showed that exercise training led to a significant decrease in fibroblasts and increase in endothelial cells in the scWAT of lower BMI participants. This finding aligns with our current understanding from rodent models, where exercise training has been shown to decrease fibrosis and increase angiogenesis^50,13,14^. Moreover, our results suggest heterogeneous cellular mechanisms underlying scWAT adaptations in distinct human participant groups. Additionally, there were significant increases of myeloid cells, fibroblasts, and ASCs, and decreases of adipocytes and vascular cells in higher BMI compared to lower BMI participants at baseline. Together, both the deconvolution data findings and the participant group differences in the transcriptomic adaptations are novel findings in human adipose tissue biology and enhance our understanding of the molecular mechanisms underlying obesity and T2D.

A particularly novel finding from this work is that circadian rhythm-related genes that were downregulated at baseline in participants with higher BMI were upregulated by exercise training, although this reversal only occurred in higher BMI participants without diabetes. Deconvolution analysis identified mature adipocytes as the primary cell type mediating this effect. Key circadian rhythm genes have been reported to modulate lipolysis, thermogenesis, oxidative stress as well as expression and secretion of adipokines in mature adipocytes *in vitro*^51^. As disruptions in the circadian rhythm have been implicated in both obesity and T2D^52,53^, exercise training may restore circadian rhythm gene misalignment in WAT functioning to help ameliorate metabolic disease.

Results showing that the extent of exercise-induced scWAT transcriptomic reprogramming correlates with changes in serum FFA indicate that adaptations to WAT may facilitate lower circulating FFAs, which in turn could lead to a reduction in systemic insulin resistance. Systolic blood pressure reduction and a trend to BMI reduction in higher BMI participants also correlated with scWAT transcriptomic adaptations. Alterations in baseline gene expression in T2D participants compared to higher BMI participants without diabetes was partially reversed with exercise training. As participants with T2D demonstrated improved glycemic control post exercise-training, an important focus for future mechanistic studies will be to determine if these select genes regulate systemic glucose metabolism. These findings suggest an important role for exercise-induced scWAT adaptations in modulating the cardiometabolic phenotype, including blood pressure regulation, glucose, and lipid metabolism in humans^50,54^ and for molecular scWAT adaptations to exercise training to reduce the detrimental effects of obesity and T2D.

In summary, these discoveries define the broad transcriptomic adaptations to scWAT that occur with obesity and T2D, and the reversal of many of these adaptations by exercise training, an intervention that has well-established metabolic health benefits. Supported by genetic evidence from GWAS studies, we speculate that specific genes may play a causal role in overall physical activity responses in people with and without metabolic disease. These data form the basis for future studies to define mechanistic roles for select genes and pathways to counter the detrimental changes of long-term obesity. Through an in-depth understanding of both the cell-type specific changes in scWAT, and TF regulation of genes and pathways, these insights on exercise-induced human scWAT adaptations will help identify novel pharmacotherapeutic targets to prevent and alleviate obesity and T2D.

### Limitations of study

This analysis was underpowered to detect male and female differences and gender-specific exercise training effects. Exercise training induces adaptations in different WAT depots, but visceral or gluteal WAT samples were not available for analysis. While training did induce changes in VO_2_peak and glycemic control, training of longer duration may exert different or additional adaptations.

## STAR□METHODS

## KEY RESOURCES TABLE

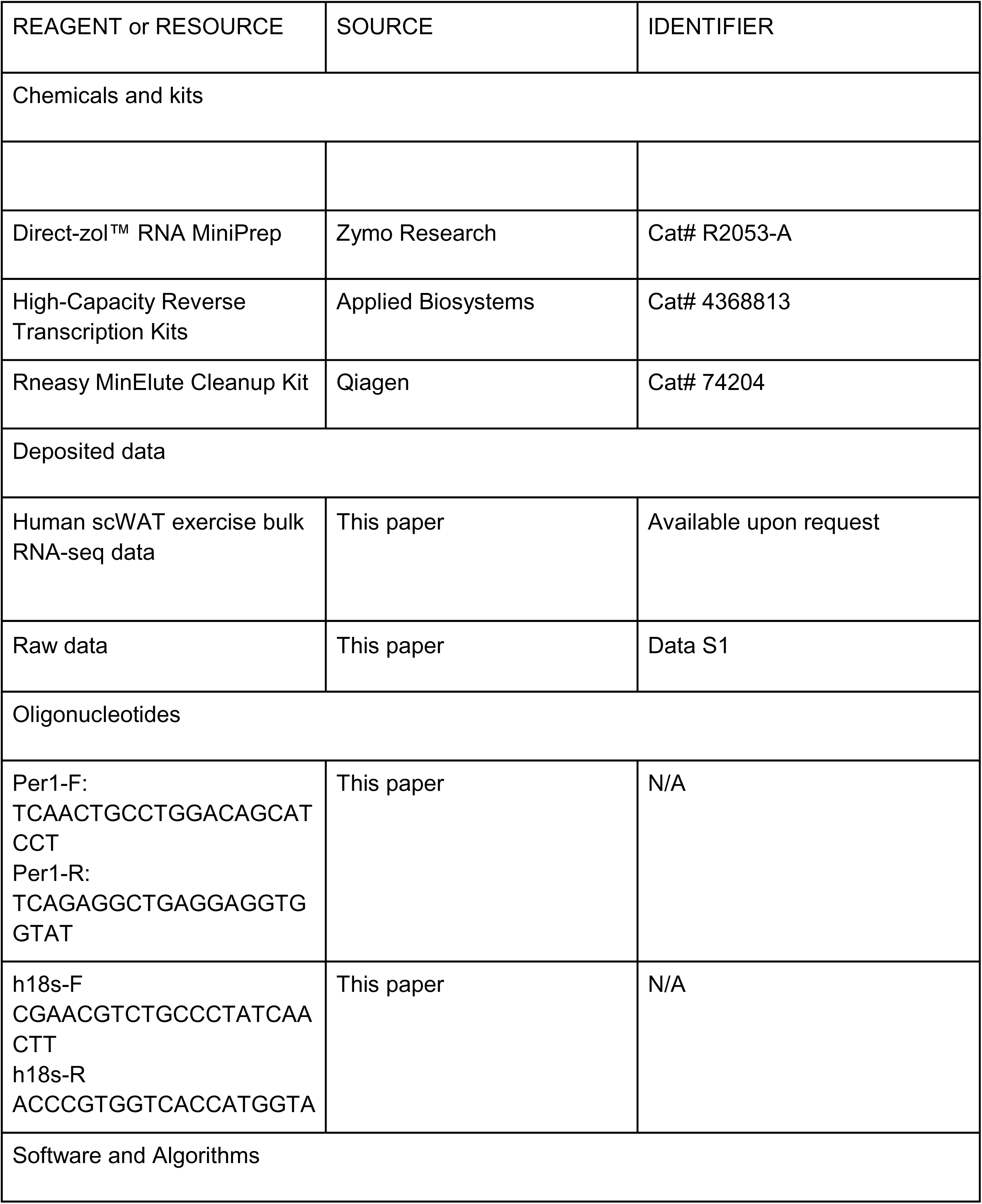
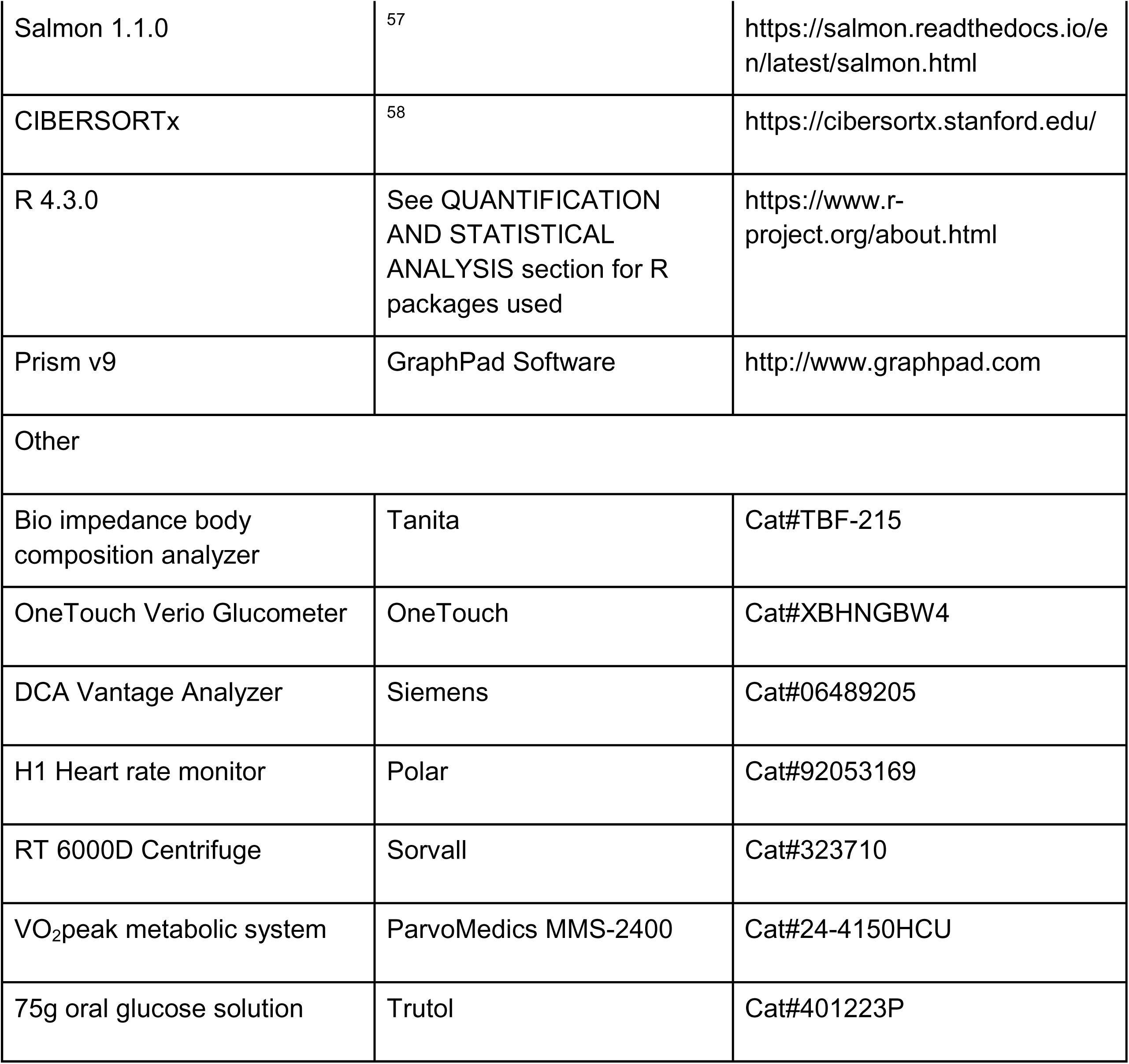

## RESOURCE AVAILABILITY

### Lead Contact

Further information and requests for resources and reagents should be directed to and will be fulfilled by the Lead Contact, Laurie J. Goodyear.

### Materials Availability

This study did not generate new unique materials. Data and Code Availability

– All raw and processed bulk RNA-seq data have been uploaded to the GEO database (https://www.ncbi.nlm.nih.gov/gds) with the accession number *available upon request*.
– Original codes for data analysis were deposited on Github at https://github.com/KellisLab/human_metab.
– All values used to generate the graphs of the paper can be found in the file Data S1 – Source Data. Any additional information required to reanalyze the data reported in this paper is available from the lead contact upon request.

### Study population

Male and female adults with or without overweight/obesity and type 2 diabetes were enrolled in a supervised 10-week moderate-intensity exercise training program. Overall study inclusion criteria included: BMI 20-37 kg/m^2^, age 25-55 years, HbA1c ≤9.0%. The clinical trial was conducted with three study arms: 1) ‘Lower BMI’ participants 20.0-26.9 kg/m^2^, 2) ‘Higher BMI’ participants without T2D, categorized by BMI 27.0-35.0 kg/m^2^, and 3) ‘T2D’ participants with higher BMI and established or newly diagnosed type 2 diabetes with HbA1c between 6.5-9% and BMI between 25.0-35.0 kg/m^2^. The analysis was conducted on the first 30 enrolled participants in the study. All participants were previously sedentary according to the American College of Sports Medicine’s guidelines^59^. Exclusion criteria consisted of: type 1 diabetes, severe complications of diabetes, heart or lung disease, current dieting or weight loss efforts, cancer, renal or hepatic dysfunction, neurological disease, clinical history of stroke, uncontrolled hypertension, and inability to exercise at 50% of predicted heart rate reserve. Participants taking beta-blockers were excluded. Participants were asked to maintain a constant diet and avoid weight-loss efforts during the study. All study-related tissue collections occurred in the morning, before noon. No participants were taking beta blockers. During the training program, participants with T2D continued their current medication regimen. Anti-hyperglycemic medications used by participants included: Biguanides (6/8), GLP-1 receptor agonists (3/8), SGLT2-inhibitors (1/8), Sulfonylureas (2/8), and Glargine insulin (1/8). All participants provided written informed consent before enrollment in the study. The consent process was conducted by ethical guidelines, ensuring that participants were fully informed of the study’s purpose, procedures, risks, and benefits. Participants were informed that their participation was voluntary and they could withdraw at any time without any penalty or loss of benefits. The study was approved by the Joslin Diabetes Center’s Institutional Review Board and registered at https://Clinicaltrials.gov (#NCT03133156).

### Body composition analysis, blood, and tissue collections

At the first and last visit, participants arrived after an overnight fast. Blood was collected for fasting glucose, insulin, serum lipids, and markers of kidney and liver function. An oral glucose tolerance test (OGTT) was performed pre-and post-training with sampling at 0, 15, 30, 60, 90, and 120 min. Capillary blood glucose was measured by glucometer (Onetouch), serum insulin was measured by ELISA (Mercodia). HbA1C was measured by Siemens DCA Vantage Analyzer or an HbA1c documented in Joslin Diabetes Center’s electronic medical record within 2 months before the start of the study was used. Free fatty acids were determined using the NEFA-Hr (2) Reagents kit (FUJIFILM Medical Systems)^60^. Body composition was obtained by bioimpedance pre- and post-training (Tanita, Arlington Heights, Illinois). Pre- and post-training, participants underwent a periumbilical subcutaneous adipose tissue biopsy using a modified liposuction technique^14^. The white adipose tissue was immediately cleaned, processed, and stored for further analyses in liquid nitrogen.

### Exercise training intervention

To determine peak aerobic capacity, a non-fasted, standard VO_2_peak test was conducted by the exercise physiologist at pre- and post-training. A ramp style protocol on the cycle ergometer was used for the VO_2_peak test^61^ and blood pressure, heart rate, and (Rating of Perceived Exertion) RPE were measured. Next, the 10-week moderate intensity continuous exercise training (MIT) protocol began, based on a target heart rate range corresponding to 70-75% of VO_2_peak using the heart rate reserve formula^59^. Initially, participants exercised 20-30 min/day at ∼40-50% VO_2_peak, 3d/week. By week 4, the duration, intensity, and frequency of exercise were gradually increased to 45-60 min/day at ∼70-75% of VO_2_peak, 4x/week, by week 4. Participants choose the mode of aerobic exercise they performed; running, biking, elliptical, rowing, or Arch trainer. At least two exercise sessions per week were completed on-site, while the additional sessions were monitored via activity tracker to assure >90% compliance. Lower BMI participants were randomized to either MIT as above, or a high-intensity interval training (HIT) training program. For HIT, the first 2 sessions consisted of 4 x 30s all-out cycling efforts with 4min of recovery in between. For session 3 and 4, 5 x 30s efforts were performed, and the remainder of the sessions were 6 x 30s efforts. The HIT training occurred three times/week on-site for 10 weeks.

### White adipose tissue RNA isolation and RNA-sequencing

Subcutaneous white adipose tissue of 30 participants (N=60 samples) was included in this analysis (as part of trial #NCT03133156). RNA was isolated according to standard procedures we have published^14^. Briefly, RNA was isolated from human subcutaneous white adipose tissue using an RNA extraction kit (Direct-zol™ RNA MiniPrep, Zymo Research). RNA was further purified with high quality using RNeasy MinElute Cleanup Kit (Qiagen). RNA was reverse-transcribed using standard reagents (High Capacity Reverse Transcription Kits, Applied Biosystems), and cDNA was amplified by RT-PCR. For each gene, mRNA expression was calculated relative to 18s. Libraries were prepared using a modified version of the Takara SMARTer Stranded Total RNA-seq Kit–Pico Input Mammalian kit. In brief, 50ng of RNA at 5 ng/μL was sonicated using R230 sonicator (Covaris), and the resulting material was confirmed using a Fragment Analyzer (Agilent). Five nanograms of each sonicated sample was prepared using the pico input kit as for FFPE samples using a 1:8 volume reduction on the STP MosquitoHV. Final libraries were validated by Fragment Analyzer and qPCR prior to sequencing on a NextSeq500 with 50 nt single-end reads.

### RNA-seq analysis of samples collected before and after exercise training

We ran Salmon 1.1.0^57^ to quantify the number of unique reads for each transcript against Ensembl version 98 human transcripts. The transcript level information was summarized to the gene-level using R package tximport^62^. We then performed principal component analysis (PCA) of the baseline samples prior to exercise training using normalized counts by the DESeq2 “vst” function^63^. Based on visual inspection of the first and second principal components (PCs), we observed sex-specific expression patterns and determined genes showing a significant sex-specific expression difference at baseline with a P-value <0.01 (Wilcoxon rank-sum test). We removed genes that had a TPM (Transcripts Per Kilobase Million) value of zero in >30% of the participants in this analysis and the downstream differential expression gene (DEG) analysis. We removed the two samples from participant 3 from further analyses because the baseline sample from this participant was an outlier on the PCA plot. After outlier sample removal, we re-performed PCA for the baseline samples to reveal a separation of the baseline samples along the lower and higher BMI axis at PC1 and the female and male axis at PC2. Sex-specific genes (**Table S1**) were labeled and interpreted with caution in the downstream DEG analysis whenever the two sexes were combined.

To visualize global transcriptomic changes of the post-training samples relative to their corresponding baseline samples, we performed matrix multiplication using the normalized counts of the post-training samples and the PCA eigenvectors from the baseline samples, and obtained the PC coordinates of the post-training samples in the same PCA space as the baseline samples. To correlate the global changes with the changes in clinical measurements, we calculated euclidean distances between the post-training and their corresponding baseline samples using the first two PCs or all the PCs, and then tested the distance differences for discrete variables (higher vs. lower BMI and female vs. male) using student’s t-test, and the distance associations with continuous variables (age, delta weight, etc.) using Pearson’s product moment correlation coefficient. We identified the clinical measurements with a nominal p-value <0.05 as significantly associated with the global transcriptomic changes in scWAT.

For DEG analyses, we used TPM values for the analyses. Wilcoxon rank-sum test was performed between groups. For baseline sample comparisons, we used unpaired tests; and to detect training effects, we used paired tests. We ran paired tests for sex-specific groups as well to detect sex-specific training effects. Genes with nominal p-values <0.05 and absolute log2 fold changes >0.5 were considered significant. Adjusted p-values were calculated using the Benjamini & Hochberg method (FDR)^64^.

The R package fgsea^25^ (version 1.20.0) was used to perform gene set enrichment analysis. 2982 Canonical Pathways gene sets downloaded from the MSigDB Collections (https://www.gsea-msigdb.org/gsea/msigdb/collections.jsp#C2) were used for the analysis and genes were ranked according to their differential expression significance and their fold change (-log10(p-value) * sign(logFC)). The eps argument (lower bound) of the fgsea function was set to zero to estimate p-values more accurately. Gene sets with P-values <0.05 were considered significant.

Gene Regulatory Network analysis was done using the R package DoRothEA^26^ (version 1.6.0). Only regulons with a confidence level A or B were selected for the analysis. The run_viper function was used to generate normalized enrichment scores for each transcription factor using arguments minsize=5 and eset.filter = FALSE.

### Deconvolution of RNA-seq data using publicly available human single-cell WAT data

Single-cell and single-nucleus RNA-seq data of human WAT cells from two resource papers^32,33^ were used as reference single-cell datasets. They are accessible from the Gene Expression Omnibus (GEO) and the ArrayExpress public databases with accession numbers GSE155960 and E-MTAB-9199, respectively. Cell clustering and cell-type annotation were done for each dataset exactly as described in the method section of the aforementioned articles. The R package Seurat^65^ (version 4.1.1) was used to perform scRNA-seq integration to correct for batch effects when merging the two single-cell datasets. Specifically, the select Integration Features function was used to select features that were repeatedly variable across the two datasets, cross-dataset pairs of cells in a matched biological state (‘anchors’) were identified using the FindIntegrationAnchors function, and finally the IntegrateData function was used to integrate the two datasets by using these anchors. Scaling, principal component analysis (PCA, npcs = 30 and resolution = 0.5) and UMAP dimensional reduction were carried out to visualize and cluster the integrated single-cell dataset. Finally, the number of cells were downsampled to 200 per cluster using the WhichCells function to achieve fast computing.

We used CIBERSORTx^58^ to impute cell-type abundances and cell-type-specific gene expression profiles from our bulk WAT gene expression data. A cell-type based signature matrix was generated from the integrated single-cell dataset described above. Cell fractions and high-resolution cell expression were inferred using the signature matrix and the bulk RNA-seq dataset. Finally, the gene subset file, specifically required for the high-resolution cell expression module, contained the name of each gene found in the bulk RNA-seq dataset. After obtaining the high-resolution cell expression profiles for each cell type in each sample, we performed similar DEG analysis as described above and used nominal p-values <0.05 without a log2 fold change threshold to identify cell-type-specific DEGs.

### Genetics Enrichment Analysis

Genetic enrichment of gene sets utilized stratified LD score regression by selecting ENCODE cCRES^66^ linked in subcutaneous adipose tissue using EpiMap linked above a score of 0.001^67^, when connected to a gene. For TF enrichment analysis, we used PLINK v2 using --tag-r2 0.8 -- show-tags to expand lists of variants related to moderate-to-vigorous physical activity during leisure time (GCST90104341). We then overlapped this with BSS01665 (subcutaneous adipose) enhancer states from EpiMap to obtain cCREs related to subcutaneous adipose tissue, and included GTEx v8 subcutaneous adipose eQTLs. Motif enrichment of DEG-linked peaks was then performed by counting JASPAR 2022 motifs and comparing to the subcutaneous adipose cCREs not selected using a hypergeometric test.

### Statistical Analysis for clinical measures and qPCR

For statistical analysis, Prism v9 (GraphPad) was used. Data are described as mean ± SD. Significance was accepted as p ≤ 0.05. Sample sizes are indicated in the figure legends. Statistical significance was analyzed by One-way ANOVA or paired two-tailed Student’s t-test.

## Supplemental Information

**Table S1.** Genes differentially expressed between male and female participants. Includes gene names, mean expression values in males and female, fold changes (males vs. females), log fold changes (males vs. females), raw and adjusted p-values (males vs. females).

**Table S2.** Differentially expressed genes (DEGs) between higher and lower BMI participants at baseline, including sex-specific results. Includes DEGs, mean expression values by BMI group, fold changes (higher vs. lower BMI), log fold changes (higher vs. lower BMI), raw and adjusted p-values (higher vs. lower BMI).

**Table S3.** Pathway enrichment analysis from fgsea using ranked full gene lists. Includes pathway names, p-values, adjusted p-values, enrichment scores (ES, NES), pathway size, and leading-edge genes.

**Table S4.** Transcription factor (TF) enrichment analysis. NES: normalized enrichment score. TF: transcription factor. Targets up: gene(s) predicted to be upregulated by transcription factor. Targets down: gene(s) predicted to be downregulated by transcription factor.

**Table S5.** Differentially expressed genes (DEGs) in response to exercise training, pre-vs. post-training by BMI group. Includes DEGs, mean expression values for pre-and post-training, fold change (pre-vs. post-training), log fold change (pre-vs. post-training), and adjusted p-values (pre-vs. post-training).

**Table S6.** Pathway enrichment analysis of exercise training-induced DEGs. Shows significantly up- and downregulated pathways across BMI and T2D groups, including immune, ECM, and metabolic pathways.

**Table S7.** Transcription factor (TF) enrichment analysis of exercise training-induced DEGs. TF: transcription factor. Targets up: gene(s) predicted to be upregulated by transcription factor. Targets down: gene(s) predicted to be downregulated by transcription factor.

## Acknowledgments

We would like to thank Jeffrey Richard, Taylor Pierce, Julianne O’Connell, and the CRC staff for their assistance. We also thank the study participants for their participation. We thank Stuart Levine for generating high-quality data and technical support on the RNAseq analysis.

This study was supported by NIH NIDDK grants: K23-DK114550 and BADERC P&F funding (to R.J.W.M), T32-DK110919 (to J.Y.), R01-DK112283 and R01-DK099511 (to L.J.G.), F32-DK126432 and Joslin Diabetes Center P&F (to M.V), and P30-DK36836 (to Joslin Diabetes Center), P30-DK057521 (to BADERC).

## Author contributions

R.J.W.M. directed the research project, designed research, carried out experiments, analyzed data, and wrote the manuscript. J.Y. oversaw the bioinformatics analysis, analyzed the data, and wrote the manuscript. P.N. designed research, carried out experiments, analyzed data, and wrote the manuscript. B.J. analyzed data, and wrote the manuscript. D.P. analyzed data. L.K.S. and N.P.C. assisted with human sample collection and study conduct. J.P. reviewed data and provided feedback on the manuscript. M.V. assisted with human study conduct, provided feedback, and edited the manuscript. L.-L.H. supervised the transcriptomics experiments. M.F.H and M.K. provided feedback and edited the manuscript. L.J.G. directed the research project, designed experiments, and wrote the manuscript. All authors reviewed and approved the final manuscript.

## Declaration of interests

R.J.W.M., M.K., and L.J.G. have received research support from Novo Nordisk, unrelated to the present study. R.J.W.M. is currently an employee and stockholder of Novo Nordisk.

