## Supplementary material for "Type 2 Diabetes and Obesity Alter Exercise Training-Induced Transcriptional Adaptations to Subcutaneous White Adipose Tissue": Figure S1

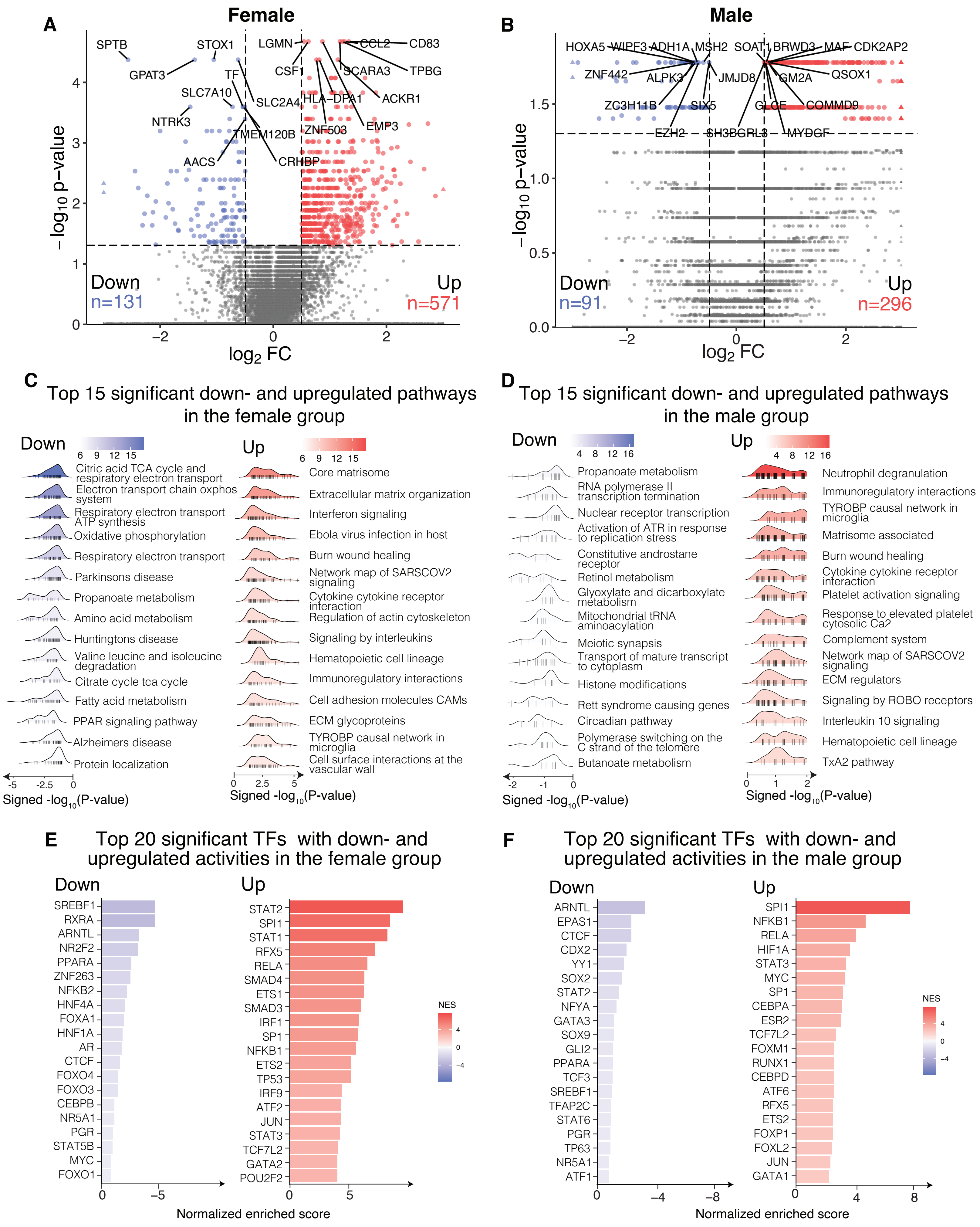
A
Female
SPTB
STOX1
GPAT3
LGMN
CCL2
CD83
CSF1
SCARA3
TPBG
TF
SLC7A10
SLC2A4
HLA−DPA1
ACKR1
NTRK3
TMEM120B
ZNF503
EMP3
AACS
CRHBP
−log10 p−value
log2 FC
Down
n=131
Up
n=571
B
Male
HOXA5
WIPF3
ADH1A
MSH2
SOAT1
BRWD3
MAF
CDK2AP2
ZNF442
ALPK3
JMJD8
GM2A
QSOX1
ZC3H11B
SIX5
GLGE
COMMD9
EZH2
SH3BGRL3
MYDGF
Down
n=91
Up
n=296
C
Top 15 significant down- and upregulated pathways in the female group
Down
Citric acid TCA cycle and respiratory electron transport
Electron transport chain oxphos system
Respiratory electron transport ATP synthesis
Oxidative phosphorylation
Respiratory electron transport
Parkinsons disease
Propanoate metabolism
Amino acid metabolism
Huntingtons disease
Valine leucine and isoleucine degradation
Citrate cycle tca cycle
Fatty acid metabolism
PPAR signaling pathway
Alzheimers disease
Protein localization
Signed -log10(P-value)
Up
Core matrisome
Extracellular matrix organization
Interferon signaling
Ebola virus infection in host
Burn wound healing
Network map of SARSCOV2 signaling
Cytokine cytokine receptor interaction
Regulation of actin cytoskeleton
Signaling by interleukins
Hematopoietic cell lineage
Immunoregulatory interactions
Cell adhesion molecules CAMs
ECM glycoproteins
TYROBP causal network in microglia
Cell surface interactions at the vascular wall
Signed -log10(P-value)
D
Top 15 significant down- and upregulated pathways in the male group
Down
Propanoate metabolism
RNA polymerase II transcription termination
Nuclear receptor transcription
Activation of ATR in response to replication stress
Constitutive androstane receptor
Retinol metabolism
Glyoxylate and dicarboxylate metabolism
Mitochondrial tRNA aminoacylation
Meiotic synapsis
Transport of mature transcript to cytoplasm
Histone modifications
Rett syndrome causing genes
Circadian pathway
Polymerase switching on the C strand of the telomere
Butanoate metabolism
Signed -log10(P-value)
Up
Neutrophil degranulation
Immunoregulatory interactions
TYROBP causal network in microglia
Matrisome associated
Burn wound healing
Cytokine cytokine receptor interaction
Platelet activation signaling
Response to elevated platelet cytosolic Ca2
Complement system
Network map of SARSCOV2 signaling
ECM regulators
Signaling by ROBO receptors
Interleukin 10 signaling
Hematopoietic cell lineage
TxA2 pathway
Signed -log10(P-value)
E
Top 20 significant TFs with down- and upregulated activities in the female group
Down
SREBF1
RXRA
ARNTL
NR2F2
PPARA
ZNF263
NFKB2
HNF4A
FOXA1
HNF1A
AR
CTCF
FOXO4
FOXO3
CEBPB
NR5A1
PGR
STAT5B
MYC
FOXO1
Up
STAT2
SPI1
STAT1
RFX5
RELA
SMAD4
ETS1
SMAD3
IRF1
SP1
NFKB1
ETS2
TP53
IRF9
ATF2
JUN
STAT3
TCF7L2
GATA2
POU2F2
NES
Normalized enriched score
F
Top 20 significant TFs with down- and upregulated activities in the male group
Down
ARNTL
EPAS1
CTCF
CDX2
YY1
SOX2
STAT2
NFYA
GATA3
SOX9
GLI2
PPARA
TCF3
SREBF1
TFAP2C
STAT6
PGR
TP63
NR5A1
ATF1
Up
SPI1
NFKB1
RELA
HIF1A
STAT3
MYC
SP1
CEBPA
ESR2
TCF7L2
FOXM1
RUNX1
CEBPD
ATF6
RFX5
ETS2
FOXP1
FOXL2
JUN
GATA1
NES
Normalized enriched score

**Figure S1. Distinct genes, pathways and transcription factors in scWAT between male and female subjects at baseline. A,** Volcano plot showing differentially expressed genes (DEGs) between higher BMI and lower BMI groups within females, with the top 10 significantly up- and downregulated genes labeled. **B,** Volcano plot showing differentially expressed genes (DEGs) between higher BMI and lower BMI groups within males, with the top 10 significantly up- and downregulated genes labeled. **C,** Ridge plot showing the top 15 significant pathways enriched by the up- and downregulated DEGs in the female group. **D,** Ridge plot showing the top 15 significant pathways enriched by the up- and downregulated DEGs in the male group. **E,** Bar plots showing the inferred transcription factors (TFs) enriched in the up- and downregulated DEGs in the female group. **F,** Bar plots showing the inferred transcription factors (TFs) enriched in the up- and downregulated DEGs in the male group.
