## Supplementary material for "Type 2 Diabetes and Obesity Alter Exercise Training-Induced Transcriptional Adaptations to Subcutaneous White Adipose Tissue": Figure S2

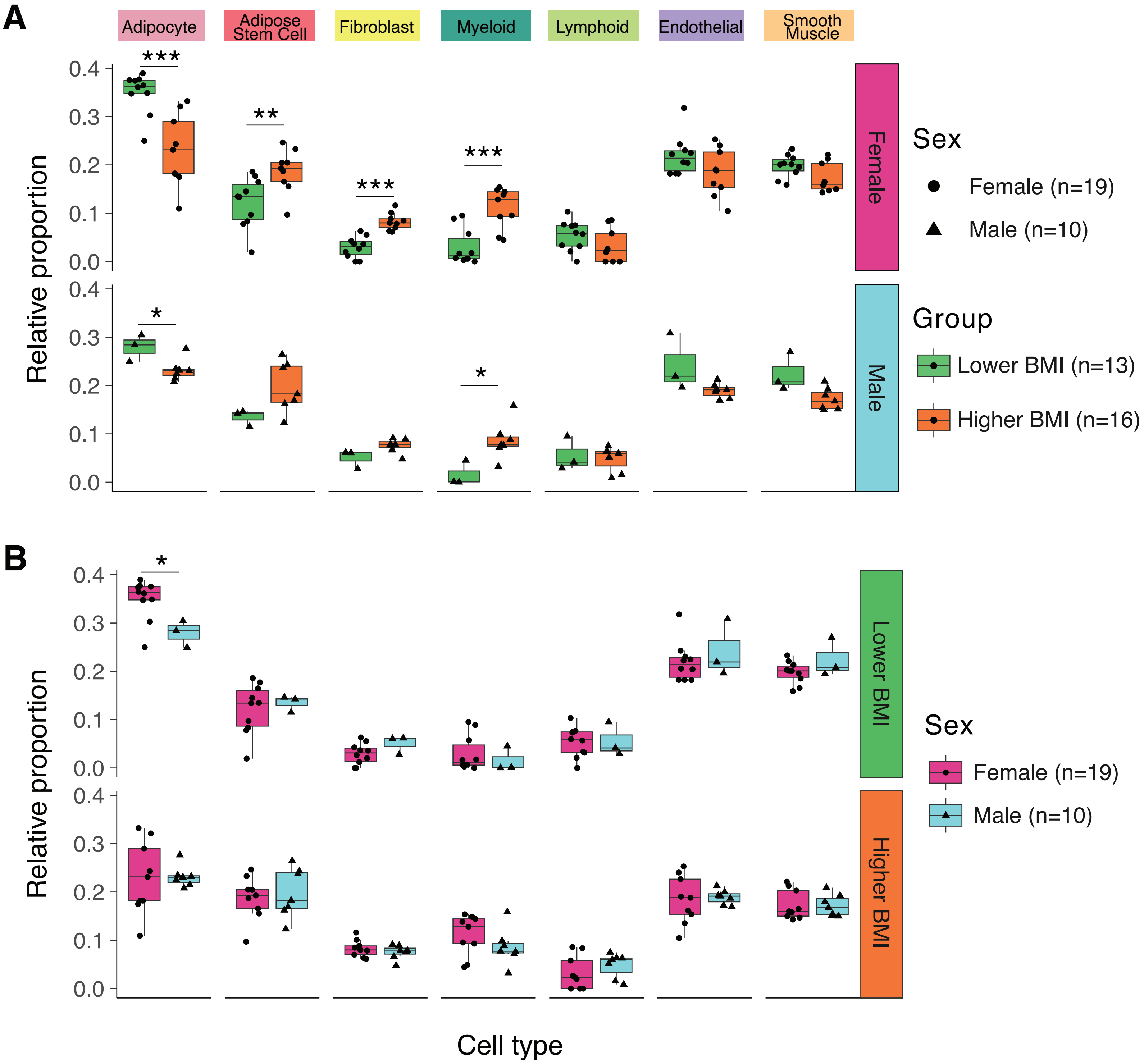


**Figure S2. Distinct cell type proportions in scWAT between male and female subjects at baseline. A,** Box plots comparing the deconvolved cell type proportions between the male and female groups by BMI category. The boxplot visualizes the median, the first and third quartiles and 1.5*inter-quartile range from the hinge. **B,** Box plots comparing the deconvolved cell type proportions between the lower and higher BMI groups by sex category. The boxplot visualizes the median, the first and third quartiles and 1.5*interquartile range from the hinge. Statistical significance was determined using the Wilcoxon Rank Sum test. *, p<0.05; **, p<0.01; ***, p<0.001; ****, p<0.0001.
