## Supplementary material for "Type 2 Diabetes and Obesity Alter Exercise Training-Induced Transcriptional Adaptations to Subcutaneous White Adipose Tissue": Figure S3

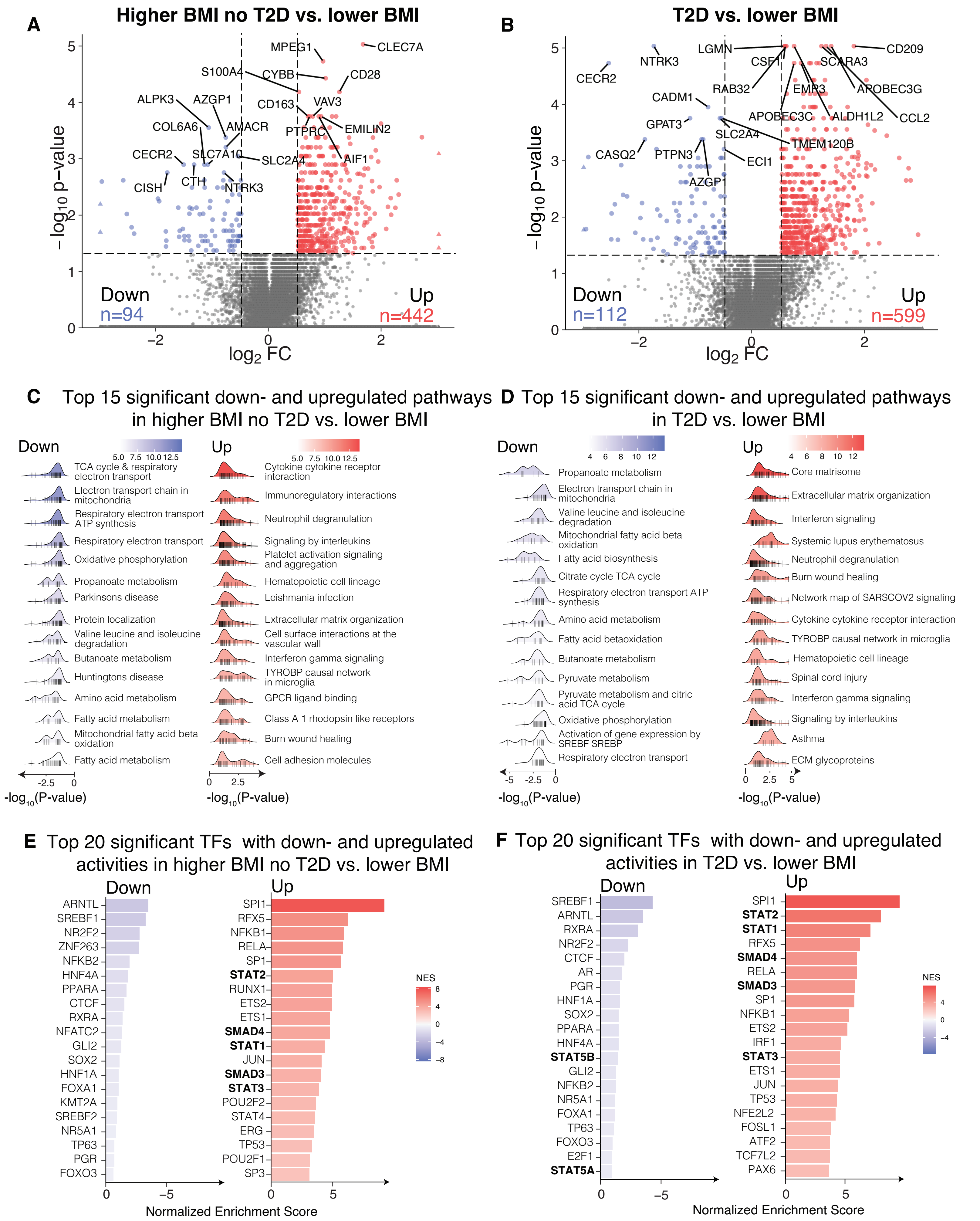
A
Higher BMI no T2D vs. lower BMI
MPEG1
CLEC7A
S100A4
CYBB
CD28
ALPK3
AZGP1
CD163
VAV3
COL6A6
AMACR
PTPRC
EMILIN2
CECR2
SLC7A10
SLC2A4
AIF1
CISH
CTH
NTRK3
Down
n=94
Up
n=442
−log10 p−value
log2 FC
B
T2D vs. lower BMI
LGMN
CD209
NTRK3
CSF1
SCARA3
CECR2
RAB32
EMR3
APOBEC3G
CADM1
APOBEC3C
ALDH1L2
CCL2
GPAT3
SLC2A4
TMEM120B
CASQ2
PTPN3
ECI1
AZGP
Down
n=112
Up
n=599
−log10 p−value
log2 FC
C Top 15 significant down- and upregulated pathways in higher BMI no T2D vs. lower BMI
Down
TCA cycle & respiratory electron transport
Electron transport chain in mitochondria
Respiratory electron transport ATP synthesis
Respiratory electron transport
Oxidative phosphorylation
Propanoate metabolism
Parkinsons disease
Protein localization
Valine leucine and isoleucine degradation
Butanoate metabolism
Huntingtons disease
Amino acid metabolism
Fatty acid metabolism
Mitochondrial fatty acid beta oxidation
Fatty acid metabolism
-log10(P-value)
Up
Cytokine cytokine receptor interaction
Immunoregulatory interactions
Neutrophil degranulation
Signaling by interleukins
Platelet activation signaling and aggregation
Hematopoietic cell lineage
Leishmania infection
Extracellular matrix organization
Cell surface interactions at the vascular wall
Interferon gamma signaling
TYROBP causal network in microglia
GPCR ligand binding
Class A 1 rhodopsin like receptors
Burn wound healing
Cell adhesion molecules
-log10(P-value)
D Top 15 significant down- and upregulated pathways in T2D vs. lower BMI
Down
Propanoate metabolism
Electron transport chain in mitochondria
Valine leucine and isoleucine degradation
Mitochondrial fatty acid beta oxidation
Fatty acid biosynthesis
Citrate cycle TCA cycle
Respiratory electron transport ATP synthesis
Amino acid metabolism
Fatty acid betaoxidation
Butanoate metabolism
Pyruvate metabolism
Pyruvate metabolism and citric acid TCA cycle
Oxidative phosphorylation
Activation of gene expression by SREBF SREBP
Respiratory electron transport
-log10(P-value)
Up
Core matrisome
Extracellular matrix organization
Interferon signaling
Systemic lupus erythematosus
Neutrophil degranulation
Burn wound healing
Network map of SARSCOV2 signaling
Cytokine cytokine receptor interaction
TYROBP causal network in microglia
Hematopoietic cell lineage
Spinal cord injury
Interferon gamma signaling
Signaling by interleukins
Asthma
ECM glycoproteins
-log10(P-value)
E Top 20 significant TFs with down- and upregulated activities in higher BMI no T2D vs. lower BMI
Down
ARNTL
SREBF1
NR2F2
ZNF263
NFKB2
HNF4A
PPARA
CTCF
RXRA
NFATC2
GLI2
SOX2
HNF1A
FOXA1
KMT2A
SREBF2
NR5A1
TP63
PGR
FOXO3
Up
SPI1
RFX5
NFKB1
RELA
SP1
STAT2
RUNX1
ETS2
ETS1
SMAD4
STAT1
JUN
SMAD3
STAT3
POU2F2
STAT4
ERG
TP53
POU2F1
SP3
NES
Normalized Enrichment Score
F Top 20 significant TFs with down- and upregulated activities in T2D vs. lower BMI
Down
SREBF1
ARNTL
RXRA
NR2F2
CTCF
AR
PGR
HNF1A
SOX2
PPARA
HNF4A
STAT5B
GLI2
NFKB2
NR5A1
FOXA1
TP63
FOXO3
E2F1
STAT5A
Up
SPI1
STAT2
STAT1
RFX5
SMAD4
RELA
SMAD3
SP1
NFKB1
ETS2
IRF1
STAT3
ETS1
JUN
TP53
NFE2L2
FOSL1
ATF2
TCF7L2
PAX6
NES
Normalized Enrichment Score

**Figure S3. Distinct genes, pathways and transcription factors in scWAT between higher and lower BMI without T2D, and between T2D and lower BMI. A,** Volcano plot showing the differentially expressed genes (DEGs) between higher BMI without T2D and lower BMI groups, with the top 10 significantly up- and downregulated genes labeled. **B,** Volcano plot showing the DEGs between the T2D group compared to lower BMI group, with the top 10 significantly up- and downregulated genes labeled **C,** Ridge plot showing the top 15 significant pathways enriched by the up- and downregulated DEGs between higher BMI without T2D and lower BMI groups. **D,** Ridge plot showing the top 15 significant pathways enriched by the up- and downregulated DEGs between T2D and lower BMI groups. **E,** Bar plots showing the inferred transcription factors (TFs) enriched in the up- and downregulated DEGs between higher BMI without T2D and lower BMI groups. **F,** Bar plots showing the inferred transcription factors (TFs) enriched in the up- and downregulated DEGs between T2D and lower BMI groups.
