## Supplementary material for "Type 2 Diabetes and Obesity Alter Exercise Training-Induced Transcriptional Adaptations to Subcutaneous White Adipose Tissue": Figure S4

A

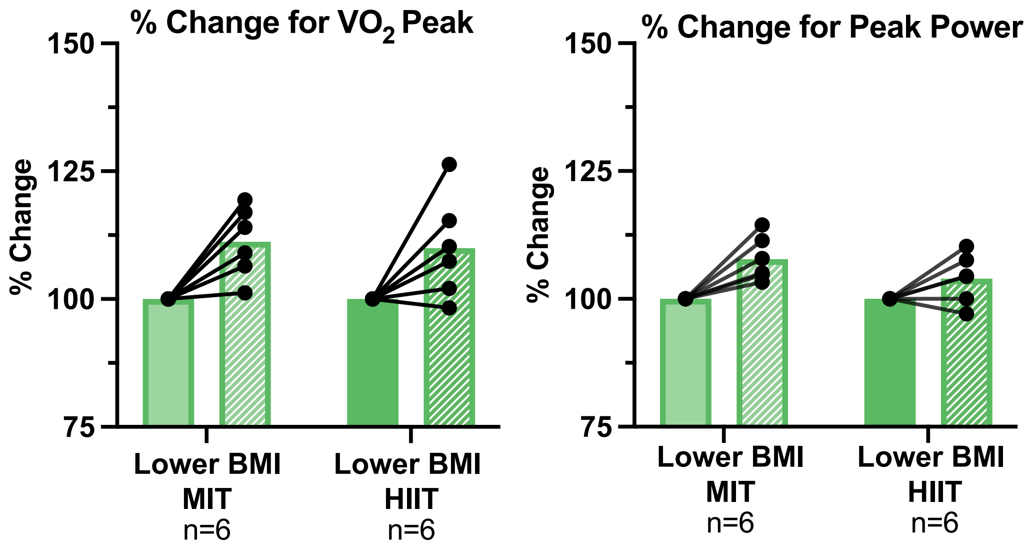


**Figure S4. Effects of training modality on cardiorespiratory fitness and BMI on transcriptomic changes in scWAT. A,** Percent change in $VO_2$ peak and peak power post-training in lower BMI moderate intensity training (MIT) and high intensity interval training (HIIT) relative to baseline.
