## Supplementary material for "Type 2 Diabetes and Obesity Alter Exercise Training-Induced Transcriptional Adaptations to Subcutaneous White Adipose Tissue": Figure S5

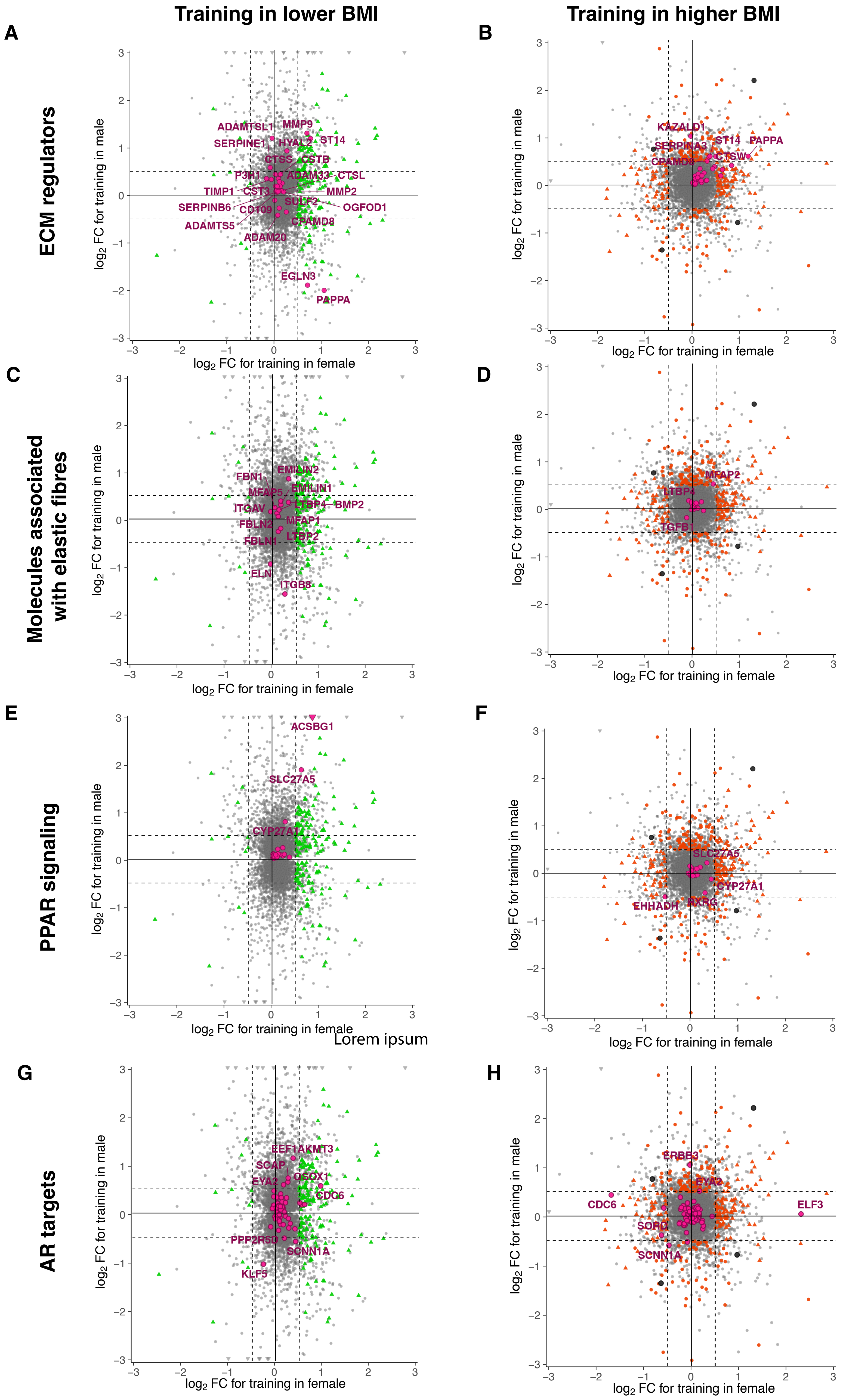
Training in lower BMI
Training in higher BMI
A
B
C
D
E
F
G
H
ECM regulators
Molecules associated with elastic fibres
PPAR signaling
AR targets
log2 FC for training in male
log2 FC for training in female
ADAMTSL1
MMP9
SERPINE1
HYAL2
ST14
CTSS
CSTB
P3H1
ADAM33
CTSL
TIMP1
CST3
MMP2
SERPINB6
CD109
SULF2
OGFOD1
ADAMTS5
CPAMD8
ADAM20
EGLN3
PAPPA
KAZALD1
SERPINA3
CTSW
FBN1
EMILIN2
MFAP5
EMILIN1
ITGAV
LTBP4
BMP2
FBLN2
MFAP1
FBLN1
LTBP2
ELN
ITGB8
MFAP2
TGFB1
ACSBG1
SLC27A5
CYP27A1
EHHADH
RXRG
Lorem ipsum
EEF1AKMT3
SCAP
QSOX1
EYA2
CDC6
PPP2R5D
SCNN1A
KLF5
ERBB3
ELF3
SORD

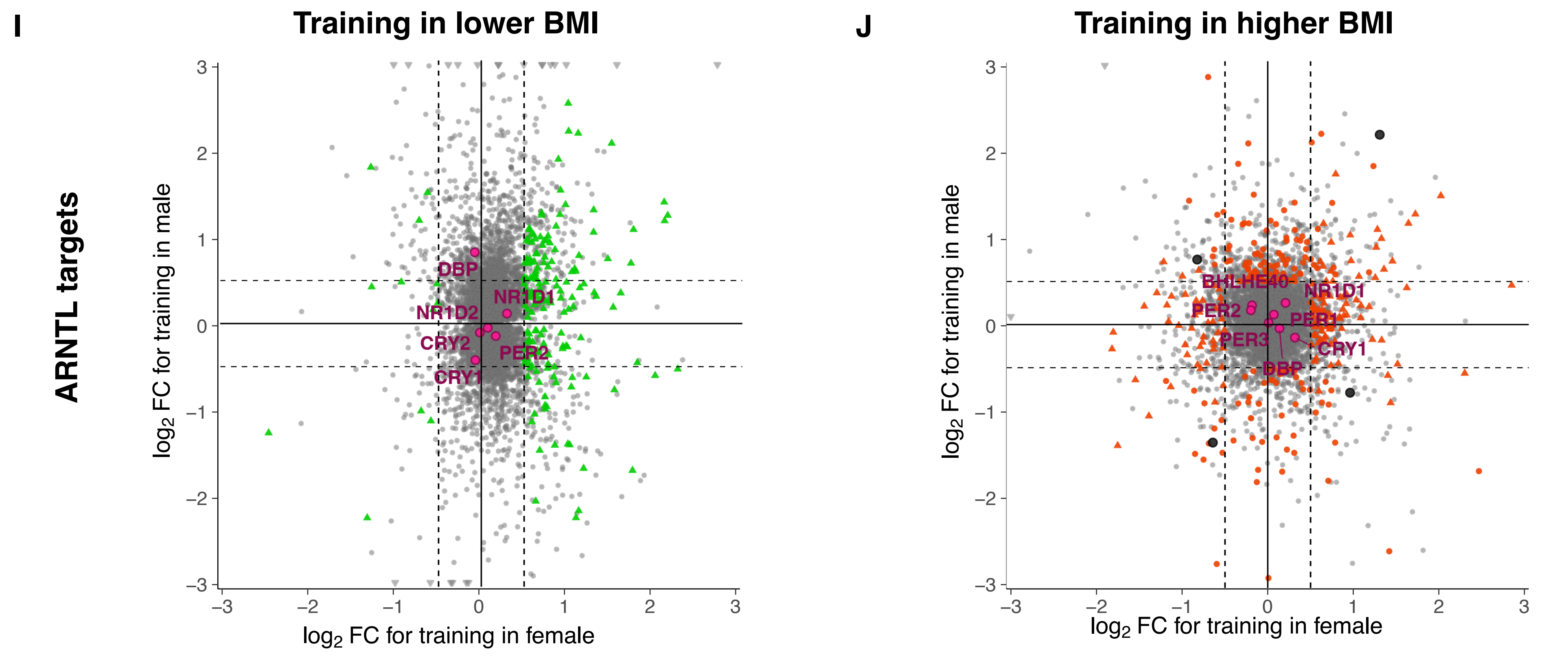


**Figure S5. Sex-specific and BMI-dependent transcriptional responses to exercise: ECM remodeling, elastic fibre dynamics, PPAR pathway regulations, AR targets, and ARNTL targets. A,B** Scatterplot comparing the training effect of gene expression in extracellular matrix (ECM) regulators in lower BMI group **(A)** and higher BMI group **(B)** in males and females. **C,D** Scatterplot comparing the training effect of gene expression in molecules associated with elastic fibresin lower BMI group **(C)** and higher BMI group **(D)** in males and females. **E,F** Scatterplot comparing the training effect of gene expression PPAR signaling in lower BMI group **(E)** and higher BMI group **(F)** in males and females. **G,H** Scatterplot comparing the training effect of gene expression in AR targets in lower BMI group **(G)** and higher BMI group **(H)** in males and females. **I,J** Scatterplot comparing the training effect of gene expression in ARNTL targets in lower BMI group **(I)** and higher BMI group **(J)** in males and females**.** Magenta indicates genes in the pathway. Green indicates significance at $p<0.05$ in lower BMI group. Orange indicates significance at $p<0.05$ in higher BMI group.
