## Supplementary material for "Type 2 Diabetes and Obesity Alter Exercise Training-Induced Transcriptional Adaptations to Subcutaneous White Adipose Tissue": Figure S6

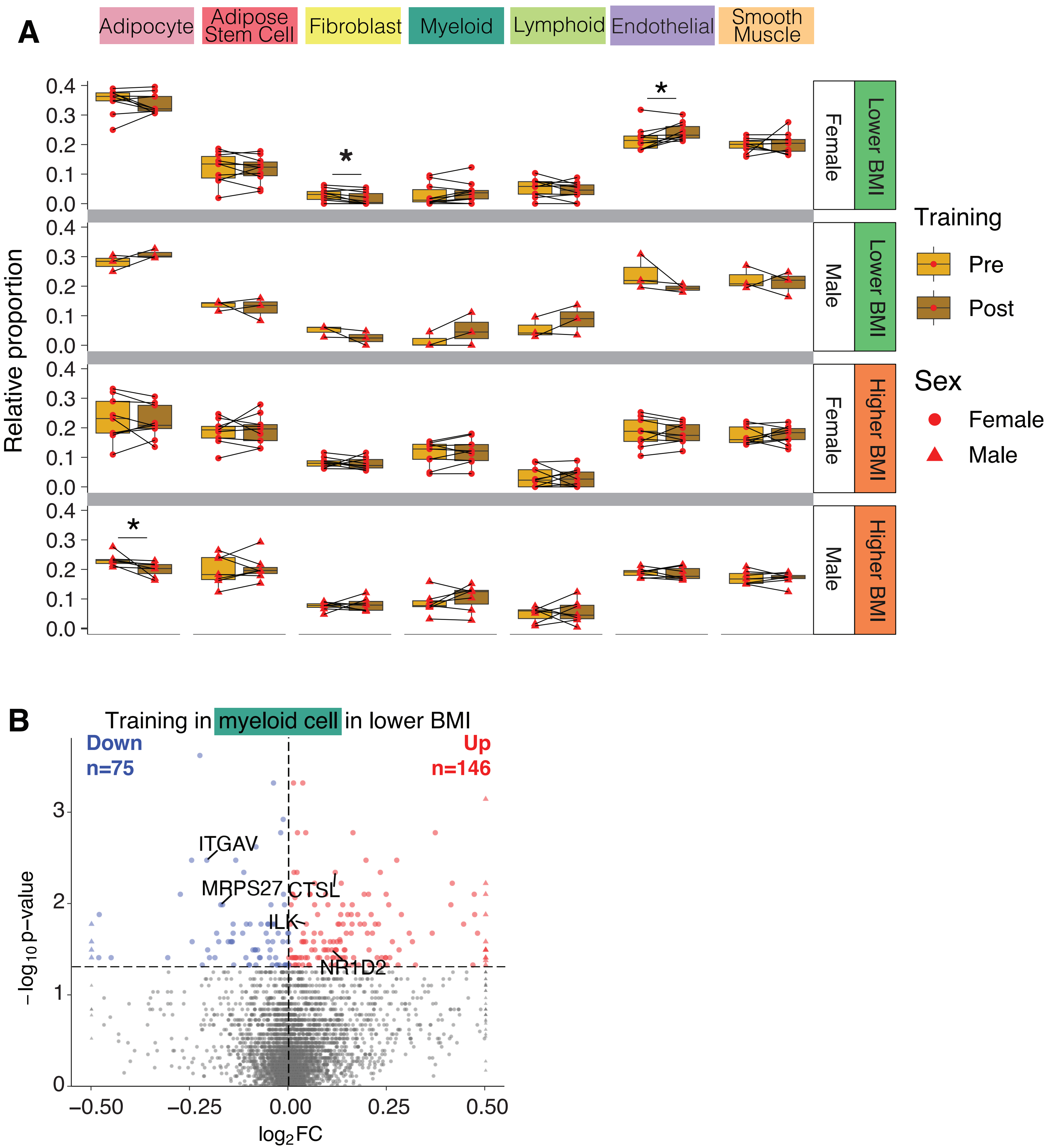


**Figure S6. Distinct cell-type proportions in scWAT before and after exercise training in lower and higher BMI groups. A,** Box plots comparing the deconvolved cell type proportions before and after exercise training for the four groups defined by BMI and sex. **B,** Volcano plot showing the exercise training effects in DEGs up- and down- regulated in myeloid cells in the lower BMI group.
