## Supplementary material for "Type 2 Diabetes and Obesity Alter Exercise Training-Induced Transcriptional Adaptations to Subcutaneous White Adipose Tissue": Figure S7

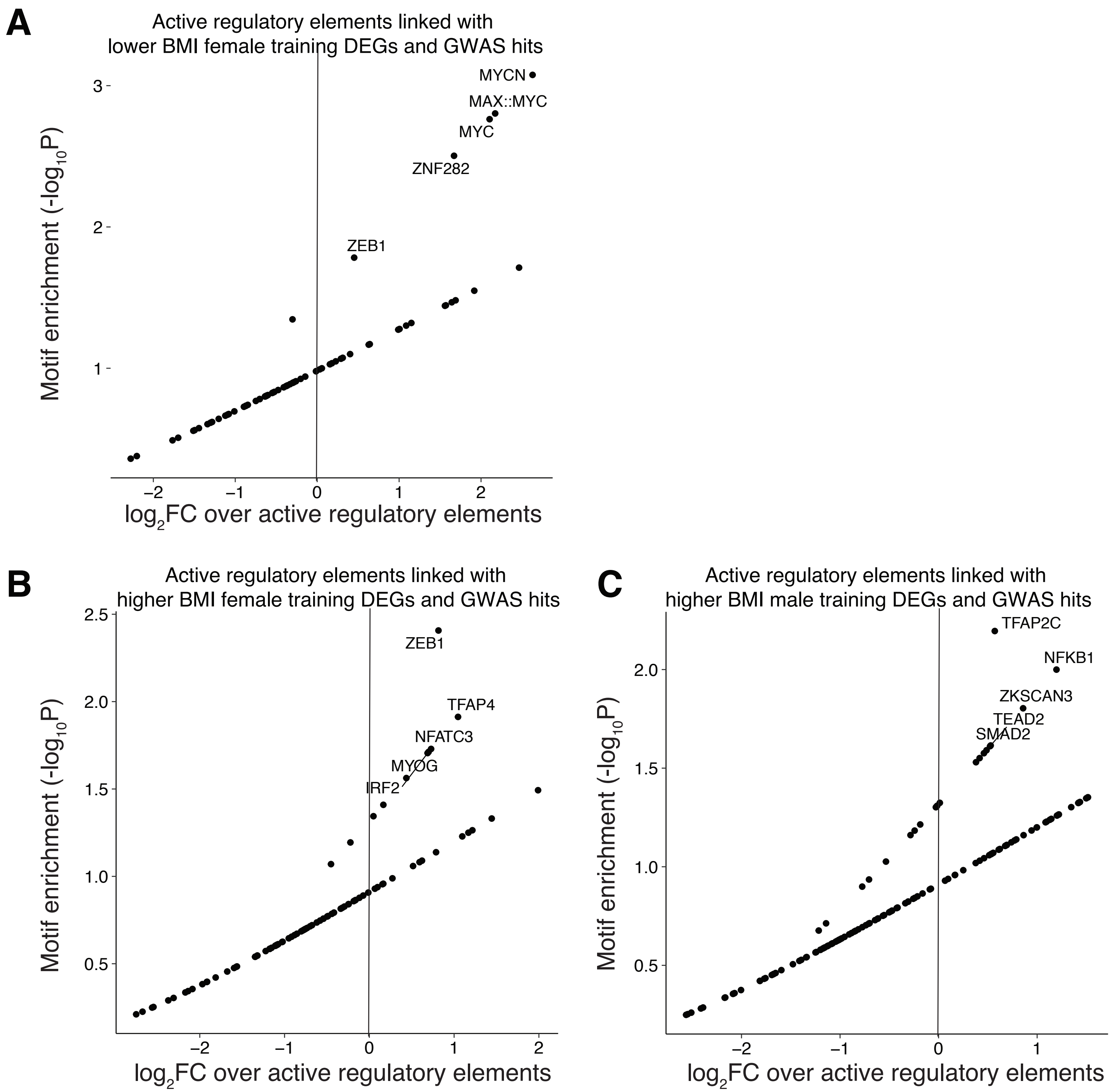


**Figure S7. Active regulatory elements associated with BMI and sex training-induced DEGsand GWAS hits. A,** Motif enrichment analysis of active regulatory elements associated with training-induced DEGs and GWAS hits in lower BMI females. **B,** Motif enrichment analysis of active regulatory elements associated with training-induced DEGs and GWAS hits in higher BMI females. **C,** Motif enrichment analysis of active regulatory elements associated with training-induced DEGs and GWAS hits in higher BMI males.
